# Crop residues in wheat-oilseed rape rotation system: a pivotal, shifting platform for microbial meetings

**DOI:** 10.1101/456178

**Authors:** Lydie Kerdraon, Marie-Hélène Balesdent, Matthieu Barret, Valérie Laval, Frédéric Suffert

**Affiliations:** UMR BIOGER, INRA, AgroParisTech, Université Paris-Saclay, 78850 Thiverval-Grignon, France; UMR IRHS, INRA, Agrocampus Ouest, Université d’Angers, 49071 Beaucouzé, France

**Author notes:** contributed equally (Valérie Laval, Frédéric Suffert).

**Keywords:** community succession, microbial diversity, oilseed rape, residue microbiota, wheat

## Abstract

Crop residues are a crucial ecological niche with a major biological impact on agricultural ecosystems. In this study we used a combined diachronic and synchronic field experiment based on wheat-oilseed rape rotations to test the hypothesis that plant is a structuring factor of microbial communities in crop residues, and that this effect decreases over time with their likely progressive degradation and colonization by other microorganisms. We characterized an entire fungal and bacterial community associated with 150 wheat and oilseed rape residue samples at a plurennial scale by metabarcoding. The impact of plant species on the residue microbiota decreased over time and our data revealed turnover, with the replacement of oligotrophs, often plant-specific genera (such as pathogens) by copiotrophs, belonging to more generalist genera. Within a single cropping season, the plant-specific genera and species were gradually replaced by taxa that are likely to originate from the soil. These changes occurred more rapidly for bacteria than for fungi, known to degrade complex compounds. Overall, our findings suggest that crop residues constitute a key fully-fledged microbial ecosystem. Taking into account this ecosystem, that has been neglected for too long, is essential, not only to improve the quantitative management of residues, the presence of which can be detrimental to crop health, but also to identify groups of beneficial micro-organisms. Our findings are of particular importance, because the wheat-oilseed rape rotation, in which no-till practices are frequent, is particularly widespread in the European arable cropping systems.

## Background

Crop residues are an essential living element of agricultural soils. Smil [1] stressed that they “should be seen not as wastes but as providers of essential environmental services, assuring the perpetuation of productive agrosystems”. When left in the field in the period between two successive crops, rather than being buried immediately, crop residues contribute to the formation of soil organic carbon, improve soil structure, prevent erosion, filter and retain water, reduce evaporation from the soil surface, and increase the diversity and activity of micro-organisms in the ground [2]. No-till practices are becoming increasingly widespread, as they take advantage of these attributes [3]. However, such practices are often considered likely to increase the risk of disease epidemics [4–6]. Indeed, several leaf-, stem-, head-, and fruit-infecting micro-organisms, classified as “residue-borne” or “stubble-borne” pathogens, are dependent on host residues for survival during the period between successive crops and for the production of inoculum for their next attack [7, 8]. The epidemiological contribution of residues as an effective source of inoculum is well-established but difficult to quantify [e.g. 9] and generalise, because the nature of survival structures depends on the biology of the species. The situation is rendered even more complex by the presence of several species reported to act as crop pathogens in plants as endophytes, without symptom development in the plant, and in the soil and plant residues as saprophytes. Taking into account the inoculum from stubble-borne pathogens and possible competition with other micro-organisms, it appears likely that the expression of a disease is the consequence of an imbalance between a potentially pathogenic species and the rest of the microbial community, rather than the consequence of the mere presence of this species [10].

Residues constitute a crucial ecological niche, not only for pathogenic species, but also for non-pathogenic and beneficial species. Residues can be viewed as both a fully-fledged matrix and a transient compartment, because they originate from the plant (temporal link), are in close contact with the soil (spatial link) and degrade over the following cropping season, at rates depending on the plant species, the cropping practices used [11], and the year (climate effect). It remains unknown whether the succession of microbial communities in residues is driven primarily by plant tissue degradation or edaphic factors [12]. Many studies have investigated the structure of the microbial communities present during the life cycle of the plant [e.g. 13–15], but few have investigated the microbiota associated with plant residues. Several ecological studies have investigated the impact of the residue compartment on the structure of soil microbial communities [2, 16–19], but not the impact of the soil compartment on structure of the residue communities. The detritusphere, defined as the part of the soil attached to residues [12, 20, 21], is the most extensive and broad hotspot of microbial life in the soil [22]. The residue compartment and the detritusphere are located in close physical proximity but are considered by microbiologists to be separate trophic and functional niches [23]. A description of the residue communities and the specific changes in these communities over time might, therefore, help agronomists to understand the impact of cropping practices on crop productivity. Fungi and bacteria play important roles in the degradation of plant tissues in debris (cellulose, hemicellulose, lignin), but the interactions between them within the microbial community remain unclear, due to the lack of information about their origins (air-borne, soil-borne or plant-borne), their individual functions and the drivers of community structure in residues.

Crop rotation induces changes in the composition of the soil microbial community and usually reduces pathogen pressure [e.g. 18]. For instance, wheat yields benefit from “break crops” such as oilseed rape or other non-host crops to break the life-cycle of wheat-specific pathogens [24]. We focused here on the wheat-oilseed rape rotation, one of the most widely used cropping systems in Europe. In 2017, the areas under bread wheat and oilseed rape in France were 5.0 million ha and 1.4 million ha [25], respectively. As oilseed rape usually recurs every three years in the rotation and is used almost systematically either directly before or directly after wheat, we estimate that this classical rotation is used on almost 4.2 million ha every year. Half the area occupied by these two crops is now grown without tillage, with at least some of the residues of the preceding crop left on the soil [26]. The issue addressed here is thus directly relevant to more than 2 million ha, or about one tenth of the total arable area in France.

In this study, we deliberately focused on crop residues as a neglected, transient, but fully-fledged half-plant/half-soil compartment without describing the soil microbial communities, considering that it has been already performed in several studies [e.g. 27, 28]. We tested the specific hypothesis that plant is a structuring factor of bacterial and fungal communities in residues, and that this effect decreases over time, as contact with the soil induce progressive colonization of residues by other microorganisms. Over the last few years, high-throughput metabarcoding has become an indispensable tool for studying the ecology of such complex microbial communities [29], partly due to the difficulties in isolating fungal and bacterial species and growing them in axenic conditions. We used this approach to describe and compare changes in the microbial community of wheat and oilseed-rape residues left on the soil surface of three cultivated fields during two cropping seasons. We investigated whether the three main determinants of the diversity of fungal and bacterial communities - plant species, cropping season, and cropping system (monoculture vs. rotation, focusing on wheat residues) - affected the microbiota of crop residues.

## Methods

### Experimental design

#### Field plots and rotations

An extensive field experiment based on a wheat (W)-oilseed rape (O) rotation cropping system was carried out during the cropping seasons of 2015-2016 and 2016-2017 at the Grignon experimental station (Yvelines, France; 48°51’N, 1°58’E). This area is characterised by an oceanic climate (temperate, with no dry season and a warm summer). A combined diachronic and synchronic strategy [30] was used to investigate the dynamics of the residue microbial communities both over a two-year period on the same plot and along a chronosequence substituting spatial differences (three plots) for time differences. A first monoculture plot (WWW) was sown with the winter wheat cultivar Soissons. This plot had been cropped with wheat since 2007 and was used in previous epidemiological studies focusing on the impact of wheat debris on the development of Septoria tritici blotch [e.g. 31, 32, 33]. Two other plots were cropped with oilseed rape cv. Alpaga and wheat cv. Soissons in rotation (OWO, adjacent to the WWW plot, and WOW, located 400 m away; Fig. 1). The size of the three plots was identical (20 m × 100 m). The OWO and WWW plots are characterized by a silty clay loam soil and plot WOW is characterized by a silty loam soil. Soil texture of the three plots is presented in Additional Table S1. The three plots were not tilled during the two cropping seasons. The wheat and oilseed rape residues were left on the soil surface after harvest. Soil was superficially disturbed to a depth of 10 cm with a disc harrow 6 weeks later (late September), leaving a large portion of residue on the surface. Crops were managed in a conventional way following local practices (nitrogen fertilization, insecticide and herbicide treatments). No fungicide was sprayed on the leaves during the study.

**Figure 1.**
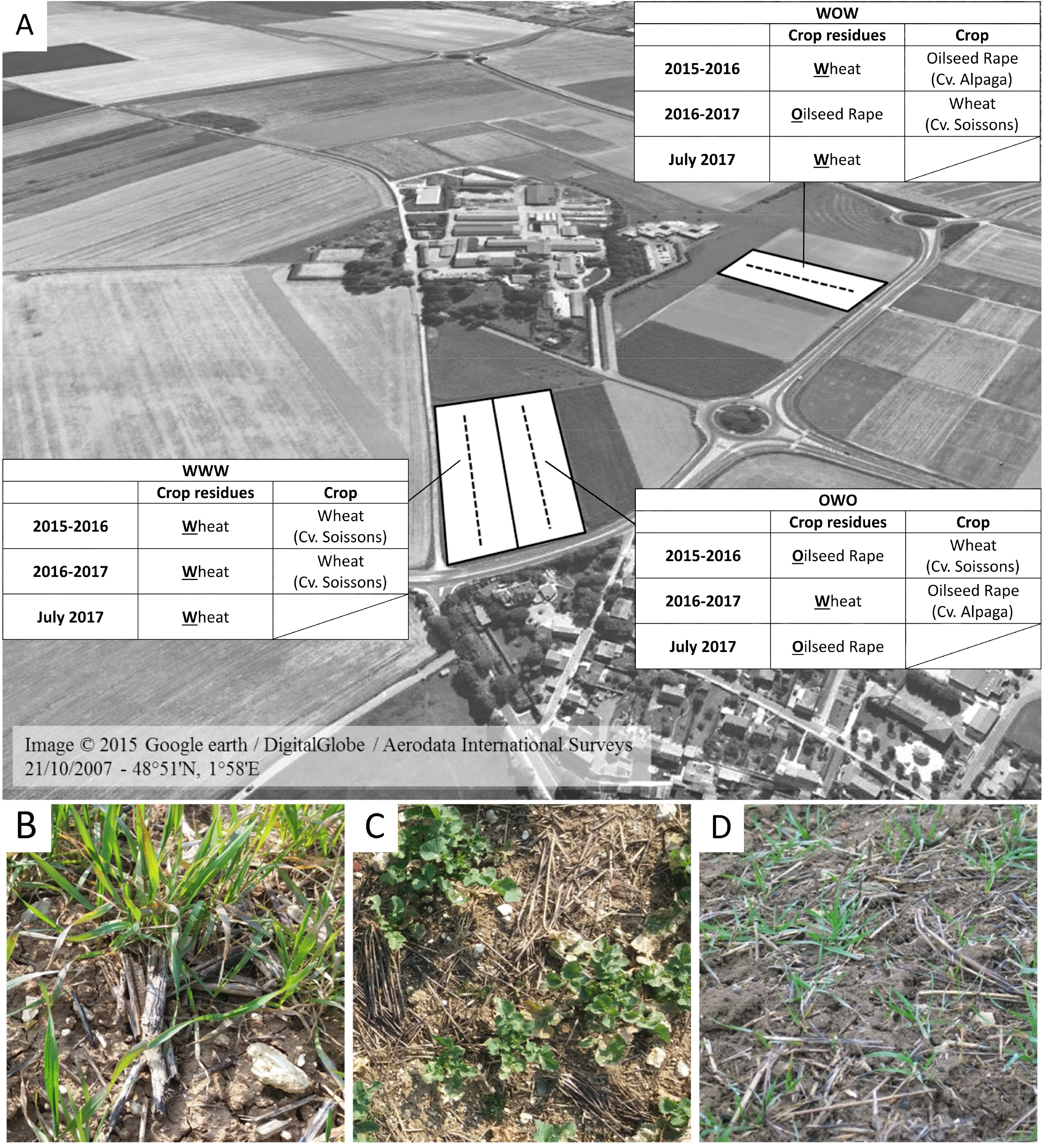
Experimental layout of the experiment. **(A)** Plots (WWW, WOW and OWO) used during the two cropping seasons of the experiment at the INRA Grignon experimental station (Yvelines, France). WWW: plot cropped with winter wheat since 2007. WOW and OWO: plots cropped with a wheat-oilseed rape rotation since 2014. Wheat straw and oilseed rape debris were chopped at harvest and a large portion of residue was then left on the surface. The dashed line indicates the sampling transect. **(B)** Oilseed rape residues in a plot cropped with wheat (OWO or WOW). **(C)** Wheat residues in a plot cropped with oilseed rape (WOW or OWO). **(D)** Wheat residues in the wheat monoculture crop (WWW).

#### Residue sampling

Wheat and oilseed rape residues from the previous crop were collected over the two cropping seasons. The changes in the microbial communities during residue degradation were described on the basis of four sampling periods each year (October, December, February, and May) Sampling dates are presented in Additional Table S2. A supplementary sample was taken in July 2016, and *a posteriori* in July 2017, to characterise the plant microbiota before the residues came into contact with the soil. For each sampling period, residues samples were collected at soil surface from five points in each plot, 20 m apart, along a linear transect (Fig. 1). Each sample was composed of twelve pieces of wheat residue or four pieces of oilseed rape residue. The five sampling points were located at the same place in the plots during the two years of the experiment.

#### DNA extraction

Residues were cut to take off remaining roots, rinsed with water to remove the soil and air-dried in laboratory conditions. They were then cut into small pieces, pooled in a 50 mL bowl and crushed with a Retsch™ Mixer Mill MM 400 for 60 seconds at 30 Hz in liquid nitrogen, in a zirconium oxide blender. The crushed powder was stored in 50 mL Falcon tubes at −80°C until DNA extraction. We transferred 40 mg of crushed residues to a 2.0 mL Eppendorf tube, which was stored to −80°C. Total environmental DNA (eDNA) was extracted according to the TriZol® Reagent protocol (Invitrogen, according to the manufacturer’s instructions). Two independent extractions were performed per sample, giving a total of 300 eDNA samples. The two extractions were considered as technical replicates.

### PCR and Illumina sequencing

Fungal and bacterial community profiles were estimated by amplifying ITS1 and the v4 region of the 16S rRNA gene, respectively. Amplifications were performed with the ITS1F/ITS2 [34] and 515f/806r [35] primers. All PCRs were run in a reaction volume of 50 µL, with 1x Qiagen Type-it Multiplex PCR Master Mix (Type-it® Microsatellite PCR kit Cat No./ID: 206243), 0.2 µM of each primer, 1x Q-solution® and 1 μL DNA (approximately 100 ng). The PCR mixture was heated at 95°C for 5 minutes and then subjected to 35 cycles of amplification [95°C (1 min), 60°C (1 min 30 s), 72°C (1 min)] and a final extension step at 72°C (10 min). PCR products were purified with Agencourt® AMPure® XP (Agencourt Bioscience Corp., Beverly, MA). A second round of amplification was performed with 5 μL of purified amplicons and primers containing the Illumina adapters and indexes. PCR mixtures were heated at 94°C for 1 min, and then subjected to 12 cycles of amplification [94°C (1 min), 55°C (1 min), 68°C (1 min)] and a final extension step at 68°C (10 min). PCR products were purified with Agencourt® AMPure® XP and quantified with Invitrogen QuantIT™ PicoGreen®. Purified amplicons were pooled in equimolar concentrations in five independent batches, and the final concentration of each batch was determined with the qPCR NGS library quantification kit (Agilent). The five independent batches were sequenced in five independent runs with MiSeq reagent kit v3 (300bp PE).

### Sequence processing

Fastq files were processed with DADA2 v1.6.0 [36], using the parameters described in the workflow for “Big Data: Paired-end” [37]. The only modification made relative to this protocol was a change in the truncLen argument according to the quality of the sequencing run. Each run was analysed separately. Taxonomic affiliations for amplicon sequence variants (ASV) generated with DADA2 were assigned with a naive Bayesian classifier on the RDP trainset 14 for bacteria [38] and the UNITE 7.1 database for fungi [39].

Only ASV detected in both technical replicates were conserved to ensure robustness [40] and were then added together. ASV classified as “Cyanobacteria/Chloroplast”, or not classified at the phylum level, were discarded from the datasets. This resulted in suppression of 1.2% of reads for fungi (4.2% of unclassified ASV), and 1.5% of reads for bacteria (4.9% of unclassified ASV and 1.3% of ASV affiliated to Cyanobacteria/Chloroplast). The remaining ASV were normalised according to the proportion of reads within each sample [41].

### Microbial community analyses

Microbial community profiles were obtained for 100 wheat residue samples and 50 oilseed rape residue samples. The alpha-diversity was estimated for each sample by calculating the Shannon index [42], completed by Faith’s phylogenetic diversity (PD; [43]; [44]) for bacterial communities. The compositional dissimilarity between samples (beta-diversity) was estimated by the Bray-Curtis dissimilarity index.

Factors taken into account in the microbial community analyses were plant (wheat; oilseed rape), cropping system (monoculture; rotation), cropping season (2015-2016; 2016- 2017), sampling period (July; October; December; February; May), and sampling plot (WWW; WOW; OWO; Fig. 1). We used a model to test these effects on each aforementioned index in a general way, and then conducted post-hoc contrasts to characterize the differences. A complete model combining all the factors could not have been used because the experimental design did not include an oilseed rape monoculture field plot. Of note oilseed rape monoculture is considered as an agronomic nonsense. Thus, a first model including the plant effect but not the cropping system effect (plant * cropping season * sampling period * sampling plot) was applied using the data set from plots in rotation only (WOW and OWO). The effect of the cropping system (monoculture; rotation) was estimated separately using a second model (cropping system * cropping season * sampling period) applied on the data set from wheat residues only; the sampling plot factor was not included in this second model as it would have been confounding with the cropping system factor.

Shannon index was calculated for both bacterial and fungal communities with the ggpubr package in R [45] and Faith’s phylogenetic diversity was calculated for bacterial communities with the picante package [46]. The effect of each factor on the Shannon index was assessed with two complementary ANOVA. A Kruskal-Wallis test also performed to assess significant differences in microbial diversity with time for each cropping season and sampling plot. Wilcoxon pairwise tests were also performed to compare the effects of sampling periods. Divergences were considered significant if *p* < 0.05.

The effect of each factor on the Bray-Curtis dissimilarity index was assessed with two complementary PERMANOVA using the adonis2 function of the vegan R package (version 2.4-4 [47] with “margin” option, used to determine the effect of each term in the model, including all other variables, to avoid sequential effects. They were visualized by multidimensional scaling (MDS) on the Bray-Curtis dissimilarity index with the phyloseq package in R (version 1.22.3 [48] and completed for bacterial communities by incorporating phylogenetic distances using the UniFrac distance matrix. After the aggregation of ASV for each sampling condition “sampling period/cropping year * crop within a rotation”, the betapart R package [49] was used to determine whether temporal changes in community composition were due to turnover (i.e. replacement of ASV between two sampling periods) or nestedness (gain or loss of ASV between two sampling periods). The effect of the plant on the microbial communities associated with residues during degradation was also assessed with PERMANOVA on each sampling period, for each cropping season.

The genus composition of fungal and bacterial communities was analysed with a cladogram based on genus names. Only genera observed in three biological samples harvested on the same plot were incorporated into the cladogram. A cladogram representing the number of ASV for each genus, read percentage, occurrence and distribution for each sample, was constructed with the Interactive Tree Of Life (iTOL [50]) online tool for phylogenetic trees.

To illustrate taxonomic changes over time, especially between plant-derived communities and communities involved later in the colonization of the residues, we focused on seasonal shifts (increase, decrease or stability) in the relative abundance of a selection of some fungal and bacterial genera and tested their statistical significance (Wilcoxon tests between sampling periods).

## Results

The bacterial and fungal communities associated with wheat (W) and oilseed rape (O) crop residues were characterised on three plots: the wheat monoculture (WWW), and two oilseed rape-wheat rotation plots (WOW and OWO) (Fig. 1). We assessed the composition of these microbial communities four times per year, during two consecutive cropping seasons (in October, December, February, and May). An additional time point (in July) was also included for identification of the micro-organisms present on the plant before contact with the soil. The analysis of raw sequence datasets for the 150 samples of wheat and oilseed rape residues collected over the two cropping seasons resulted in the grouping of 14,287,970 bacterial and 9,898,487 fungal reads into 2,726 bacterial and 1,189 fungal amplicon sequence variants (ASV). ASV not detected in both technical replicates (5.4% of bacterial reads and 1.5% of fungal reads) were removed from the datasets. Total number of reads remaining after ASV filtering is presented in Additional Table S3.

### Alpha diversity of microbial communities

Diversity dynamics, assessed by calculating the Shannon index, differed between the two cropping seasons and between fungi and bacteria. The diversity was significantly impacted by most of factors, including cropping system (monoculture; rotation) for wheat residues (Table 1; Fig. 2). Oilseed rape residues supported less fungal diversity and as much bacterial diversity than wheat residues in rotation. In addition, the diversity was significantly higher in wheat grown in monoculture than in wheat grown in rotation for both bacteria and fungi.

**Figure 2.**
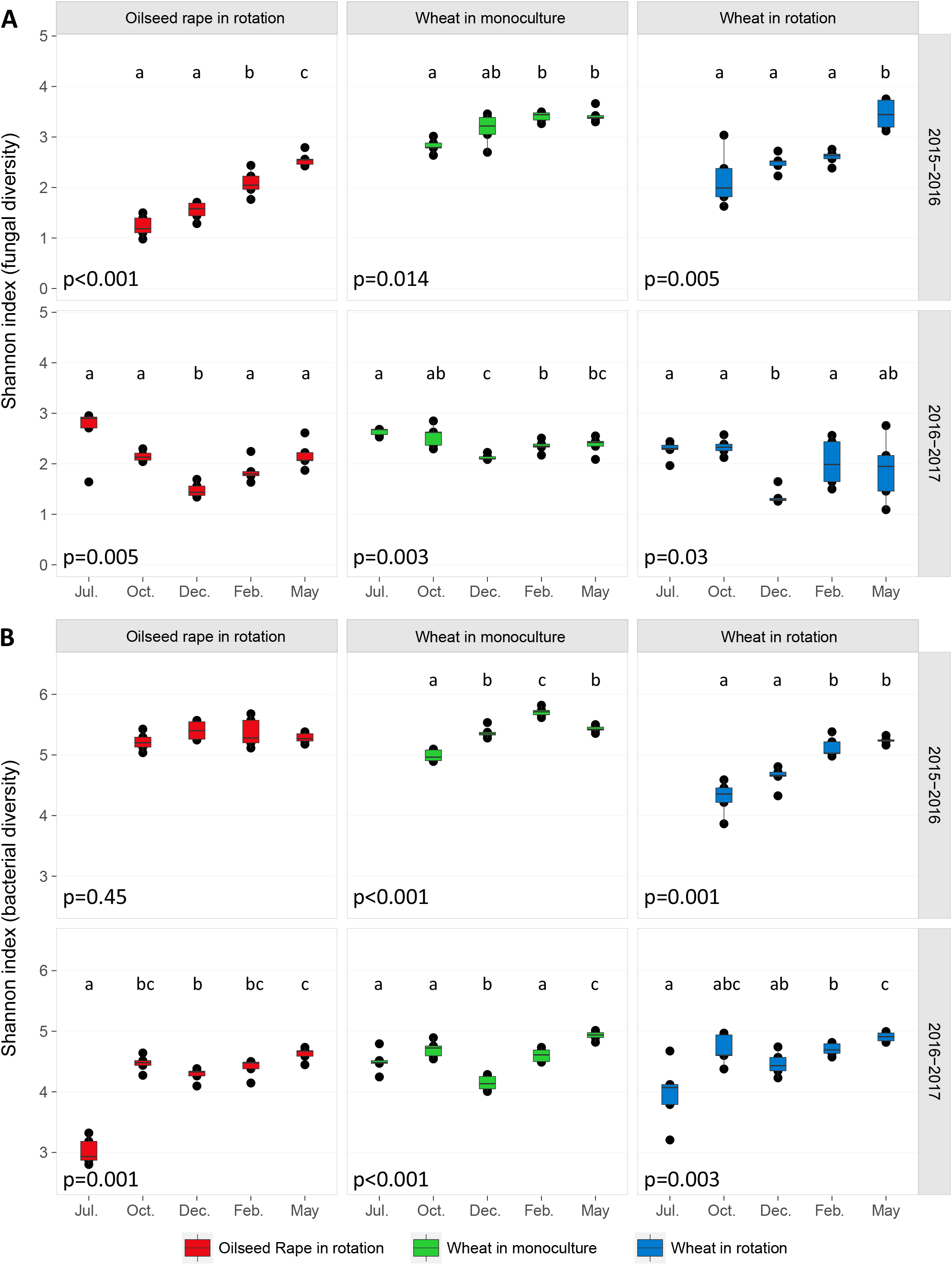
Fungal (**A**) and bacterial (**B**) diversity in plants (July) and residues (October; December; February; May), as assessed with the Shannon index, according to sampling period, the crop within a rotation (oilseed rape in WOW and OWO; wheat in WWW; wheat in WOW and OWO) and the cropping season (2015-2016; 2016-2017). Each box represents the distribution of Shannon index for five sampling points. Kruskal-Wallis tests were performed for each “crop within a rotation * cropping season” combination (*p*-values are given under each graph). Wilcoxon tests between sampling periods were performed when the Kruskal-Wallis test revealed significant differences. Samples not sharing letters are significantly different.

**Table 1.** Results of the ANOVA performed to assess the effects of plant (wheat; oilseed rape), cropping season (2015-2016; 2016-2017), sampling period (July, October; December; February; May) and sampling plot on the Shannon index of the fungal and bacterial communities present in oilseed rape and wheat residues from the plots in rotation (OWO; WOW). The effect of the cropping system (monoculture; rotation) was estimated separately with a second ANOVA performed on the wheat residue samples dataset only (wheat in monoculture, i.e. in WWW; wheat in rotation, i.e. in WOW and in OWO).

|  |  | Fungi |  | Bacteria |  |
| --- | --- | --- | --- | --- | --- |
| Tested factors |  | F value | *p*-value | F value | *p*-value |
| Plots in rotation | Plant | 22.42 | < 0.001 | 0.33 | 0.567 |
|  | Cropping season | 14.19 | < 0.001 | 35.28 | < 0.001 |
|  | Sampling period | 14.25 | < 0.001 | 37.73 | < 0.001 |
|  | Sampling plot | 31.06 | < 0.001 | 61.57 | < 0.001 |
| Wheat residues | Cropping system | 46.42 | < 0.001 | 23.66 | < 0.001 |
|  | Cropping season | 51.43 | < 0.001 | 33.34 | < 0.001 |
|  | Sampling period | 6.72 | < 0.001 | 13.89 | < 0.001 |

Fungal diversity increased over time in 2015-2016, whereas the differences between the samples in 2016-2017 did not reflect a gradual increase as the minimum was reached in December. Bacterial diversity followed a quite similar trend with however a significant decreased from February to May, more pronounced for wheat residues than for oilseed rape residues during the first cropping season; this trend was also illustrated by the Faith’s phylogenetic diversity (Additional Figure S1). The climatic conditions during residue degradation (Additional Table S4) or differences in initial diversity on the plant before harvest may explain the less marked trends observed between the two cropping seasons.

### Comparison of microbial communities associated with residues (beta diversity)

We analysed the effects of plant, cropping system, cropping season, and sampling period on communities using the Bray-Curtis dissimilarity index and PERMANOVA. There was remarkably little heterogeneity between the five samples collected in the same sampling plots (Fig. 3) and the number of biological samples was, therefore, sufficient to assess differences due to the variables of interest (i.e. plant cropping season, sampling period, and cropping system). This result was confirmed by the structure of bacterial communities visualized by incorporating phylogenetic distances using the UniFrac distance matrix (Additional Fig. S2).

**Figure 3.**
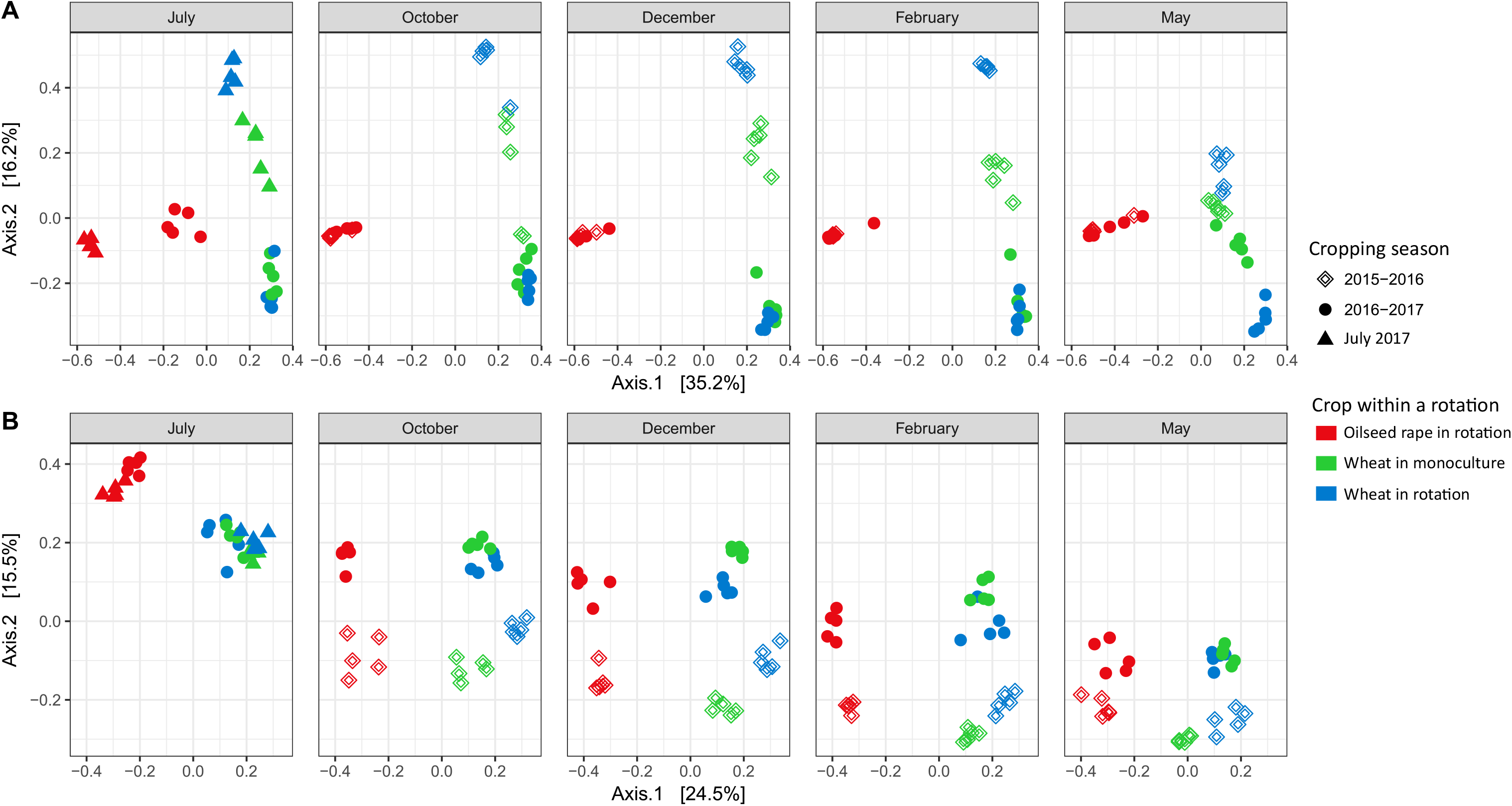
Structure of the fungal (**A**) and bacterial (**B**) communities present in oilseed rape and wheat residues, according to compositional dissimilarity (Bray-Curtis dissimilarity index), after multidimensional scaling (MDS). The two MDS were performed on the overall dataset and faceted according to the sampling period. Each point represents one sample corresponding to a cropping season (shape: 2015-2016; 2016-2017; 2017-2018) and crop within a rotation (colour: oilseed rape in rotation, i.e. in WOW and OWO; wheat monoculture, i.e. in WWW; wheat in rotation, i.e. in WOW and OWO).

#### The structure of bacterial and fungal communities is influenced by plant species and cropping system

Oilseed rape and wheat residues presented different sets of ASV, for both bacterial and fungal communities (Fig. 3). Plant species was the main factor explaining differences between the communities, accounting for 38.4% for fungi and 26.6% of the variance for bacteria, as established with PERMANOVA (Table 2). For wheat, the cropping system (rotation; monoculture) accounted for 10.5% of the variance for fungal community composition and 6.6% of the variance for bacterial community composition.

**Table 2.** Results of the PERMANOVA performed to assess the effects of plant (wheat; oilseed rape), cropping season (2015-2016; 2016-2017), sampling period (July; October; December; February; May) and sampling plot on the Bray-Curtis dissimilarity index of the fungal and bacterial communities present in oilseed rape and wheat residues from the plots in rotation (OWO; WOW). The effect of the cropping system (monoculture; rotation) was estimated separately with a second PERMANOVA performed on the wheat residue samples data set only (wheat in monoculture, i.e. in WWW; wheat in rotation, i.e. in WOW and in OWO). PERMANOVAs were performed using the adonis2 function with “margin” option.

| Data set | Factors | Fungi |  | Bacteria |  |
| --- | --- | --- | --- | --- | --- |
|  |  | R<sup>2</sup> | *p*-value | R<sup>2</sup> | *p*-value |
| Plots in rotation | Plant | 0.384 | < 0.001 | 0.266 | < 0.001 |
|  | Cropping season | 0.131 | < 0.001 | 0.118 | < 0.001 |
|  | Sampling period | 0.068 | < 0.001 | 0.149 | < 0.001 |
|  | Sampling plot | 0.100 | < 0.001 | 0.053 | < 0.001 |
| Wheat residues | Cropping system | 0.105 | < 0.001 | 0.066 | < 0.001 |
|  | Cropping season | 0.292 | < 0.001 | 0.248 | < 0.001 |
|  | Sampling period | 0.123 | < 0.001 | 0.204 | < 0.001 |

The percentage of variance explained by the plant decreased over time for fungal community structure (e.g. from 75% in October 2016 to 40% in May), while for bacteria the decrease of the percentage of variance explained by the plant was less pronounced (from 65% to 50%). The percentages of variance associated with the plant for each date are presented in Additional Table S5.

#### Community structures change over time

Cropping season was the main temporal factor underlying changes in community structure, accounting for 11.8% of the variance for bacteria when considering only the plots in rotation (or 24.8% when considering only the wheat residues without sampling plot effect), and 13.1% of the variance for fungi when considering only the plots in rotation (or 29.2% when considering the wheat residues without sampling plot effect; Table 2). Sampling period had also a significant impact on community composition, accounting for 14.9% of the variance for bacteria when considering only the plots in rotation (or 20.4% when considering only the wheat residues without sampling plot effect), and 6.8% of the variance for fungi when considering only the plots in rotation (or 12.3% when considering the wheat residues without sampling plot effect). Theoretically, changes in ASV composition result from turnover (replacement of ASV between two sampling periods) and nestedness (gain or loss of ASV between two sampling periods [49]). We found that the dissimilarity between sampling periods was smaller for bacterial than for fungal ASV structure. By decomposing the dissimilarity between sampling periods, we found that, for fungi, 94% (± 5%) of dissimilarity was explained by turnover for oilseed rape, 89% (±15%) for wheat in monoculture, and 80% (±13%) for wheat in rotation. For bacteria, 69% (±17%) of dissimilarity was explained by turnover for oilseed rape, 61% (±19%) for wheat in monoculture, and 80% (±16%) for wheat in rotation. Decomposition of dissimilarity between sampling periods is presented in Additional Table S6.

### Changes in communities, by genus

We characterised potential taxonomic differences in communities over time by analysing wheat and oilseed rape residues separately. ASV were aggregated together at genus level, resulting in 84 fungal and 184 bacterial genera for wheat, and 63 fungal and 186 bacterial genera for oilseed rape. For the sake of clarity, most 60 genera of fungi and bacteria were presented in Fig. 4 and Fig. 5, respectively. All the detected genera and their evolution over time were presented in Additional Fig. S3, S4, S5 and S6. For both plant species, we identified genera that disappeared or displayed a significant decrease in relative abundance over time. The seasonal shifts of some genera and their significance are presented in Additional Fig. S6. Among these genera, some are known to be associated with plants, such as *Alternaria*, *Acremonium* [14, 51, 52], *Cryptococcus* [53], *Sarocladium* [54] and *Cladosporium* [13, 51–54].

**Figure 4.**
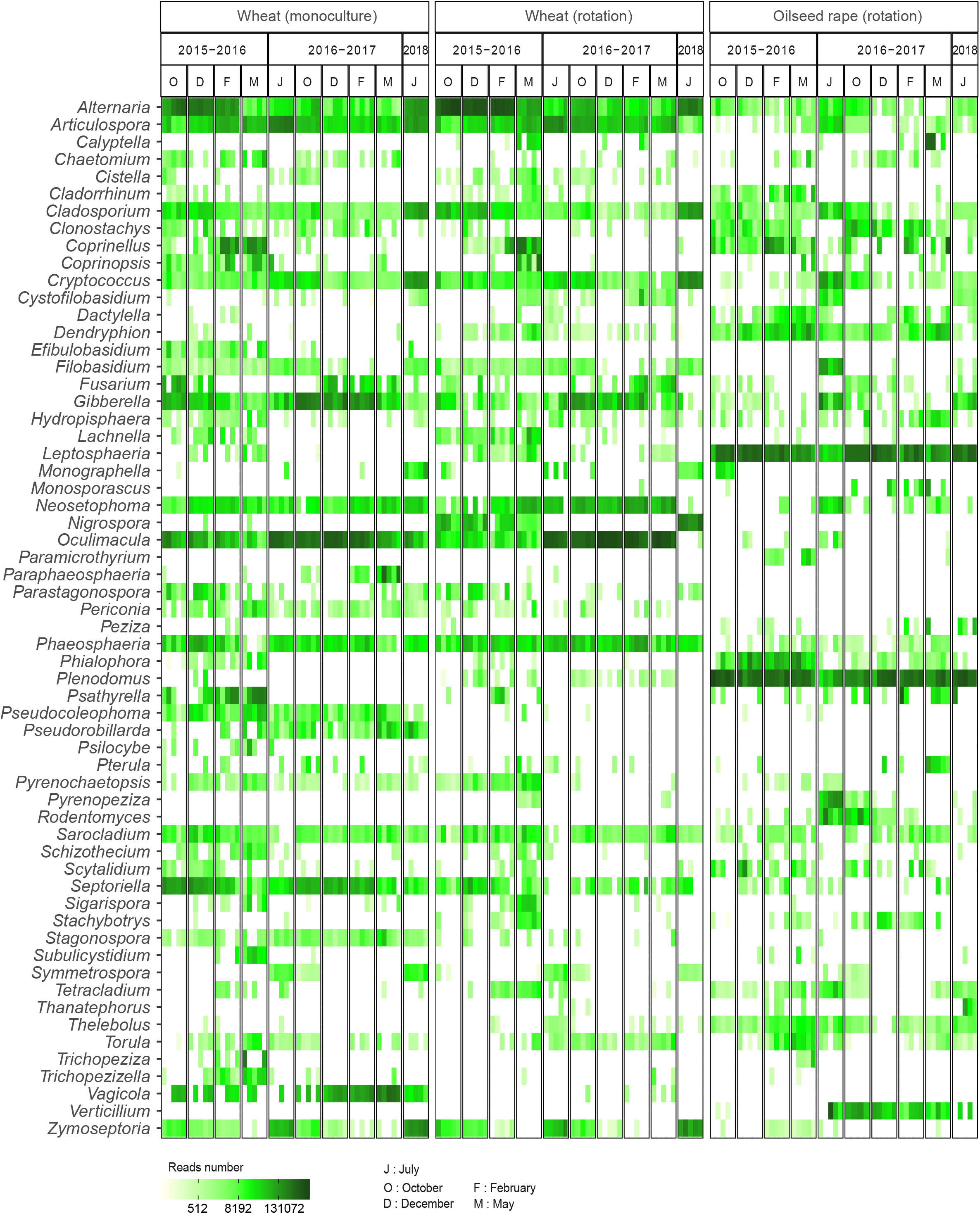
Distribution of the 60 most prevalent fungal genera detected in wheat residues in the five samples for each sampling date. Unclassified genera were removed from the visualisation.

**Figure 5.**
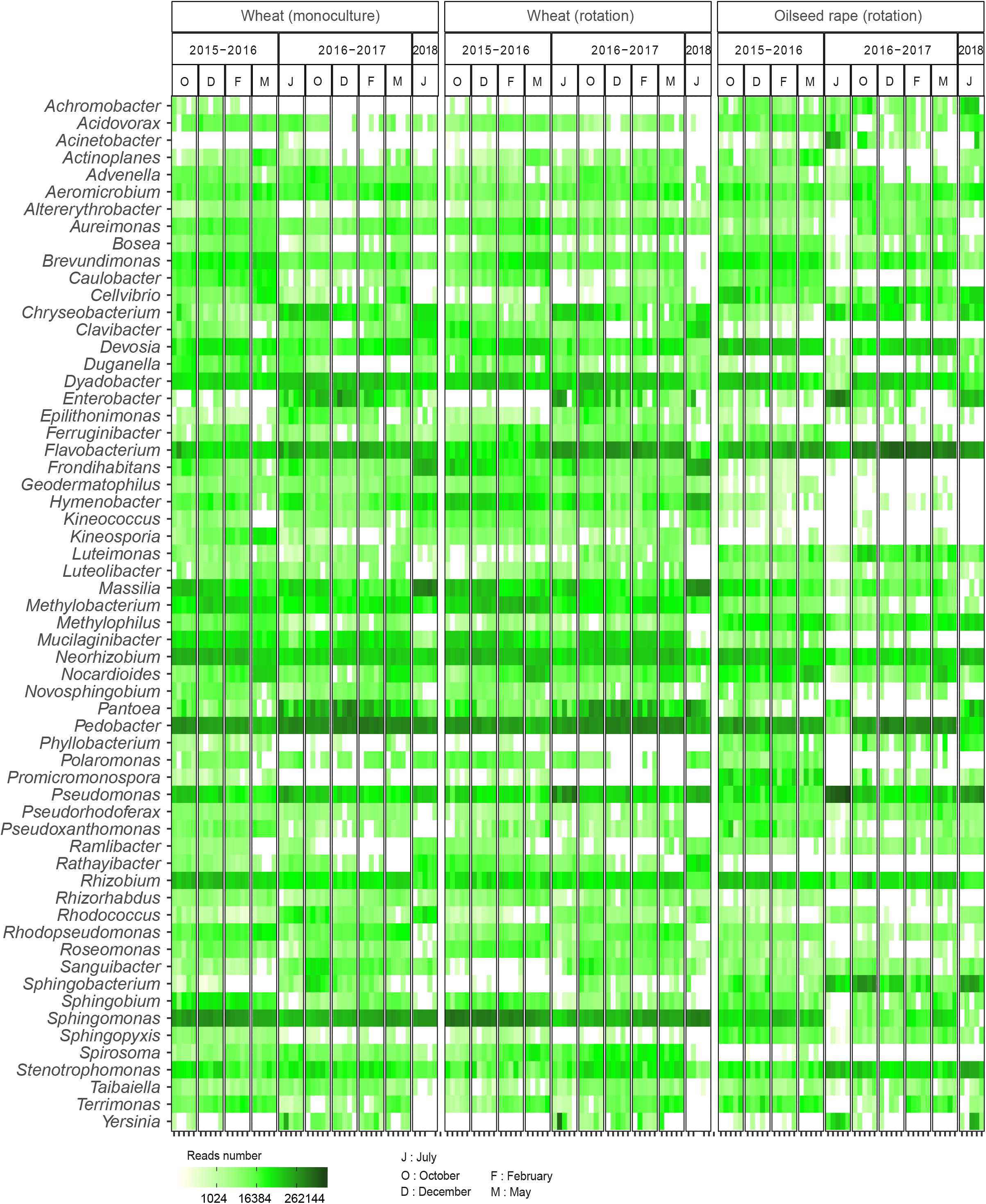
Distribution of the 60 most prevalent bacterial genera detected in wheat residues in the five samples for each sampling date. Unclassified genera were removed from the visualisation.

Some of the fungal species detected on wheat, such as *Oculimacula yallundae* (all ASV of *Oculimacula* genera), *Zymoseptoria tritici* and *Pyrenophora tritici-repentis*, (Fig. 4, Additional Fig. S3) are known to be pathogenic. Some of the species detected on oilseed rape, such as *Verticillium* spp., *Leptosphaeria maculans* (= *Plenodomus maculans*) and *Leptosphaeria biglobosa* (= *Plenodomus biglobosa*), are also known to be pathogenic. Strikingly, *L. maculans* and *L. biglobosa* predominated over the other taxa. *Verticillium longisporum*, *V. dahlia* and *V. albo-atrum* were mostly detected during the second sampling year (Fig. 4, Additional Fig. S4). As samples were collected in two different fields, it was not possible to determine whether the occurrence of *Verticillium* spp., a soil-borne pathogen complex causing *Verticillium* wilt [55], was affected more by year or by the soil contamination. *Acremonium*, *Clonostachys* and *Alternaria* genera, which have also been described as associated with plants [56], were detected in the early sampling periods. Their relative abundances decreased over time (Additional Fig. S7). Most of the genera that were not present at early sampling points and with relative abundances increasing over time (e.g. *Coprinellus*, *Psathyrella*, *Torula*, *Tetracladium*, and *Exophiala*) were common to wheat and oilseed rape residues (Fig. 4). These genera can thus be considered as probably derived primarily from the surrounding soil.

For bacteria, the difference in the genera detected between the two plants species was less marked than for fungi, as 146 genera were common to wheat and oilseed rape residues (Fig. 5). These 146 genera corresponded to the 98.7% most prevalent reads for wheat and 97.5% most prevalent genus reads for oilseed rape. *Proteobacteria* was the predominant phylum the first year. The most prevalent proteobacterial subgroup was *Alphaproteobacteria*, with a high prevalence of *Rhizobiales* and *Sphingomonadales*. *Rhizobium* and *Neorhizobium*, two major genera from *Rhizobiales*, decreased in abundance between October and May in both wheat and oilseed rape (Additional Fig. S7). *Sphingomonadales* genera were much more abundant on wheat than on oilseed rape, especially *Sphingomonas* (Fig. 5). *Bacteroidetes* genera, including *Pedobacter* in particular, were frequently detected and their prevalences tended to be stable for oilseed rape residues, and to decrease for wheat residues (Additional Fig. S7). In parallel, an increase in *Actinobacteria*, particularly *Nocarioides*, was observed. Major differences between July and October were observed for oilseed rape, consistent with the beta-diversity analysis, in which the percentage dissimilarity between July and October was high, due to both species extinction and turnover. *Gammaproteobacteria* were highly abundant on oilseed rape in July. Their frequency then decreased rapidly from October to May, due largely to the decrease in *Pseudomonas* (Fig. 5, Additional Fig. S7). In parallel, we observed an increase in the levels of *Alphaproteobacteria*, especially *Rhizobium* and *Sphingomonas*, between July and October. A small decrease in levels of *Gammaproteobacteria* was observed between July and October for wheat in rotation, whereas the percentage of reads associated with this class increased between July and December for wheat in monoculture, due largely to the decrease in *Pantoea* and *Enterobacteria* (Fig. 5, Additional Fig. S7). The abundance of *Bacteroidetes*, especially *Pedobacter* and *Flavobacterium*, also increased between July and October.

## Discussion

Most studies on crop residues have focused on their impact on soil microbial communities [16], and the rare studies investigating the impact of soil on residue communities focused exclusively on bacteria [27, 28] or fungi [57]. Most of these studies were conducted on residues from a single year. Bastian et al. [12] established an extensive description of the species present in the soil, detritusphere and wheat residues, using sterilised residues and soil in a microcosm. In this study, we showed, under natural conditions, that three main factors (plant species, cropping season, rotation) simultaneously influence the composition of both fungal and bacterial communities present on residues. This study is the first to investigate the total fungal and bacterial communities associated with wheat and oilseed rape residues by a metabarcoding approach over two consecutive years. The very low variability of the communities for the five replicates is remarkable and shows that our strategy would be appropriate for comparing the effects of different treatments on microbial communities.

### Crop residues should be viewed as a shifting platform for microbial meeting strongly affected by plant species

Oilseed rape and wheat residues contained different sets of micro-organisms before soil contact and during the firsts sampling periods after harvest. Similar results were previously obtained for the bacterial communities of buried crop residues [28]. Consistent with the findings of this previous study, the divergence between wheat and oilseed rape bacterial communities was probably due to differences in the chemical compounds present in the plants. The rapid change in the community observed at early stages of residue degradation for oilseed rape may be explained by the modification of simple compounds (sugars, starch, etc.), whereas wheat is composed of more complex compounds (lignin) and is, therefore, broken down less quickly, resulting in a slower change in the microbial community [28]. Overall, the change in bacterial community composition highlights turnover between copiotrophs and oligotrophs. Although copiotrophy and oligotrophy are physiological traits, several attempts have been made to classify microorganisms as oligotrophs and copiotrophs based on phylogeny [58]. According to this generalization, bacterial and fungal taxa whose relative abundances are significantly decreased during succession belong mainly to copiotroph. These taxa include for instance *Alternaria*, *Cladosporium*, *Massilia* and *Pseudomonas* (Additional file: Fig. S5). In contrast, the relative abundances of oligotrophic taxa such as *Coprinellus* or *Nocardiodes* increased during residues degradation, which could be indicative of the superior abilities of these micro-organisms to degrade complex polymers.

The initial fungal communities were structured mostly by the presence of species originating from the plant, several of which were highly specialised on the host plant. These species were gradually replaced by more generalist species, which colonised the residues of both plants. Most of these generalists, such as *Exophiala*, *Coprinellus* and *Torula*, are known to be soil-born [59, 60], or involved in degradation, such as *Coprionopsis* [61]. The host-specific fungi identified in our study included a large number of ascomycetes known to be foliar pathogens (*O. yallundae, Fusarium* sp. and *Gibberella* sp*., Z. tritici, P. tritici-repentis, Parastagonospora nodorum, Monographella nivalis, L. biglobosa* and *L. maculans*). The lifestyles of some pathogens are well-documented, as for *Z. tritici, P. tritici-repentis* and *L. maculans.* The decrease with time in levels of *Z. tritici* and other pathogens in wheat residues contrasts with the persistence of *L. maculans* and *L. biglobosa* in oilseed rape residues. These three pathogens are all known to reproduce sexually on the residues of their host plant [31, 62], but the life cycle of *L. maculans* is characterised by systemic host colonisation through intracellular growth in xylem vessels [63], whereas the development of *Z. tritici* is localised and exclusively extracellular [64]. Oilseed rape residues thus provide *L. maculans* with greater protection than is provided to *Z. tritici* by wheat residues. This likely explains differences in the persistence of the two pathogens and in the temporal dynamics of ascospore release: over up to two years for *L. maculans* [65, 66] but only a few months for *Z. tritici* [31, 67]. The predominance of *L. maculans* on oilseed rape residues was not surprising given that the oilseed rape cultivar Alpaga is known to be susceptible to *L. maculans*, but the high abundance of *L. biglobosa* was much more remarkable. One surprising finding of our study was the constant association of *L. maculans* with *L. biglobosa* on residues. Indeed, *L. biglobosa* is known to be more associated with upper-stem lesions [68], and its presence in large amounts on residues has never before been reported.

Our findings are consistent with current epidemiological knowledge of emblematic wheat and oilseed rape diseases, but they highlight our lack of knowledge concerning the lifestyles of many other fungal pathogens present on residues. A key point to be taken into account is that the trophic status of many species known to be principally pathogenic or non-pathogenic is not definitive [69]. For instance, *Alternaria infectoria* is sometimes described as a pathogen of wheat [13, 70], sometimes as an endophyte [71], and has even been tested as a potential biocontrol agent against *Fusarium pseudograminearum* on wheat [72]. Crop residues, half-plant/half-soil, should be the focus of future studies aiming to disentangle the succession of microbial species with different lifestyles and to characterise their relative impacts on the development of currently minor, but potentially threatening diseases.

### The residue microbiota should be analysed in a dynamic manner, both within and between years

The results of our study highlight the importance of conducting multi-year studies focusing on ecological dynamics both within and between years in natural conditions. Year had a strong effect on both bacterial and fungal communities. Fluctuations of climatic conditions (temperature, rainfall, wind) have a major impact on pathogenesis (disease triangle concept [73]) and on the saprophytic survival of plant pathogens during interepidemic periods [74]. The two years of our study were marked by similar means of 10-day mean temperatures, but large differences in rainfall: mean 10-day cumulative rainfall in the first year was almost twice that in the second (Additional Table S5). The colonisation of residues by late colonisers may be affected by such climatic differences: in wheat, most prevalent degrading fungi (like *Coprinellus*, *Psathyrella*, *Coprinopsis*) were almost absent in the second year of the study. There was also considerable dissimilarity between the bacterial communities associated with each of the two years. For example, genus *Enterobacter*, which was highly abundant in the second year, was barely detectable in the first year.

### Crop rotation has little impact on residue microbial communities

Oilseed rape is never grown in monoculture, so the effect of crop rotation was assessed only for wheat. The effect of rotation on residue microbial communities was much smaller than the effect of year (cropping season). It was more marked for fungi, for which diversity was greater in monoculture than in rotation. The use of a rotation may prevent the most strongly specialised species, in this case fungi, from becoming established, regardless of their pathogenicity. This finding is consistent with the greater development of some diseases in monoculture conditions, which promote the maintenance of pathogens through the local presence of primary inoculum. For instance, the presence of *P. tritici-repentis*, agent of tan spot disease, in the wheat monoculture plot and its absence from wheat-oilseed rape plots is consistent with epidemiological knowledge indicating that this disease can be controlled by leaving a sufficient interval between consecutive wheat crops in the same field [75].

### Lesson to be learned from the residue microbial communities for the sustainable management of debris-borne diseases: a delicate balance between pathogenic and beneficial micro-organisms

The maintenance of crop residues at the surface of the cultivated soil increases the microbial diversity of the soil and, in some ways, helps to maintain good functional homeostasis [76]. However, conservation practices tend to increase the risk of foliar diseases [4–6]. Most disease management strategies focus on epidemic periods, during which the pathogen and its host are in direct contact. Interepidemic periods are also crucial for pathogen development, although during these periods the primary inoculum is not directly in contact with the new crop whilst not present in the field. Indeed, by carrying the sexual reproduction of several fungal pathogens, residues contribute to the generation and transmission of new virulent isolates potentially overcoming resistance genes, during monocyclic epidemics, as described for oilseed rape canker caused by *L. maculans* [77], but also polycyclic epidemics, as described for Septoria tritici blotch caused by *Z. tritici* [78].

However, the results of our study suggest that residues should not only be considered as a substrate for pathogens and a potential source of inoculum. Indeed, we detected several fungi identified as beneficial or even biocontrol agents in previous studies, such as *Clonostachys rosea*, *Aureobasidium pullulans*, *Chaetomium globosum* and *Cryptococcus* spp. *C. rosea*, which was detected in both oilseed rape and wheat residues, has been reported to limit the sexual and asexual reproduction of *Didymella rabiei* on chickpea residues by mycoparasitism [79]. It has also been reported to be effective against *Fusarium culmorum* on wheat plants, through antibiosis during the epidemic period [80], and on wheat residues, through antagonism during the interepidemic period [81]. *Cladosporium* sp., which were abundant in our study, have also been reported to inhibit the development of *P. tritici*-*repentis* on wheat plants [82] and of *Fusarium* sp. on wheat residues [81]. The presence of these fungal species on wheat and oilseed rape residues is of potential interest for future analyses of interactions. Due to the use of a low-resolution marker for bacterial characterisation, we were unable to identify similarly the bacteria potentially interacting with pathogenic fungi. For instance, the presence of *Pseudomonas* spp. suggests possible interactions both with other microbial species and with the host plant [83], but the nature of the potential interactions is indeterminate: species of the *Pseudomonas fluorescens* group are known to be beneficial to plants, whereas *Pseudomonas syringae* and *Pseudomonas aeruginosa* are known to be pathogens of plants and even humans.

Although our study reveals the presence of genera or species reported in the literature as biocontrol agents, it has not yet shown any interaction between them and the pathogens. This experimental study (sampling effort, residue treatments, etc.) was not designed to characterize such interactions. A strategy involving the inference of microbial interaction networks from metabarcoding datasets might help to identify the species beneficial against pathogens, through competition, antagonism or parasitism. This however requires a more analytical, comparative experimental approach, that goes beyond the only description of shifts in natural communities composition: for example, using different “treatments” in a broad sense (e.g. artificial inoculation with a species or a group of species, change of biotic or abiotic environmental conditions, etc.) in order to modify interaction networks and so highlight the impact of some groups of micro-organisms on the whole community or a given species.

## Conclusion

This study shows that crop residues, which can be seen as half-plant/half-soil transient compartment, constitute a pivotal fully-fledged microbial ecosystem that has received much less attention than the phyllosphere and rhizosphere to date. This study therefore fills a gap in knowledge of the communities present on crop residues under natural conditions. It confirms that the microbiote of crop residues should be taken into account in the management of residue-borne diseases. Taking into account this ecosystem is essential, not only to improve the quantitative management of crop residues, but also to identify groups of beneficial micro-organisms naturally present. The beneficial elements of the microbial community should be preserved, or even selected, characterised and used as biological control agents against the pathogens that complete their life cycle on the residues. These results are particularly important in that wheat-oilseed rape rotations are among the most widespread arable cropping systems in France and Europe.

## Acknowledgements

This study was performed in collaboration with the GeT core facility, Toulouse, France (http://get.genotoul.fr) and was supported by *France Génomique National Infrastructure*, funded as part of “*Investissement d’avenir*” program managed by *Agence Nationale pour la Recherche* (contract ANR-10-INBS-09). We thank Martial Briand (INRA, UMR IRHS) and Dr. Gautier Richard (INRA, UMR IGEPP) for assistance with bioinformatic analyses, and Dr. Thierry Rouxel (INRA, UMR BIOGER) for improving and clarifying this manuscript. We thank Julie Sappa for her help correcting our English. We finally thank the reviewers for their insightful comments on the paper, as these comments led us to improve the correctness of several analyses.

## Availability of data and materials

The raw sequencing data is available from the European Nucleotide Archive (ENA) under the study accession PRJEB27255 (Sample SAMEA4723701 to SAMEA4724326). We provide the command-line script for data analysis and all necessary input files as Additional File 2.

## Authors’ contributions

LK, FS, VL, MHB, MB conceived the study, participated in its design, and wrote the manuscript. LK conducted the experiments and analysed the data. FS and VL supervised the project. All authors read and approved the final manuscript.

## Ethics approval and consent to participate

Not applicable

## Consent for publication

Not applicable

## Additional files

**Figure S1.**
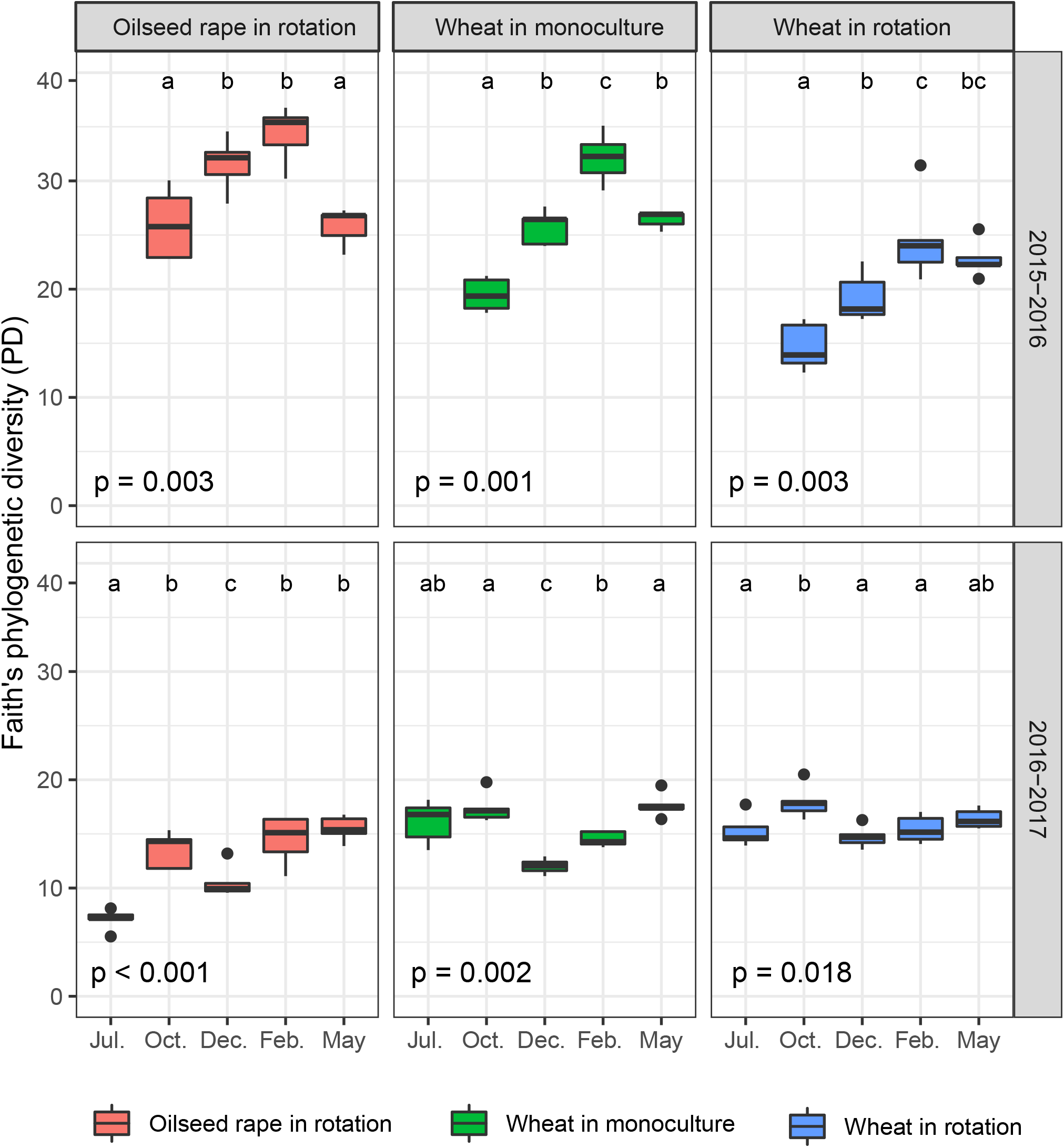
Bacterial diversity in plants (July) and residues (October, December, February, May), as assessed with the Faith’s Phylogenetic Diversity index (PD), according to sampling period, the crop within a rotation (oilseed rape in OWO or WOW, wheat in WWW, wheat in WOW or OWO) and the cropping season (2015-2016, 2016-2017). Each box represents the distribution of PD for five sampling points. Kruskal-Wallis tests were performed for each “crop within a rotation * cropping season” combination (p-values are given under each graph). Wilcoxon tests between sampling periods were performed when the Kruskal-Wallis test revealed significant differences. Samples not sharing letters are significantly different.

**Figure S2.**
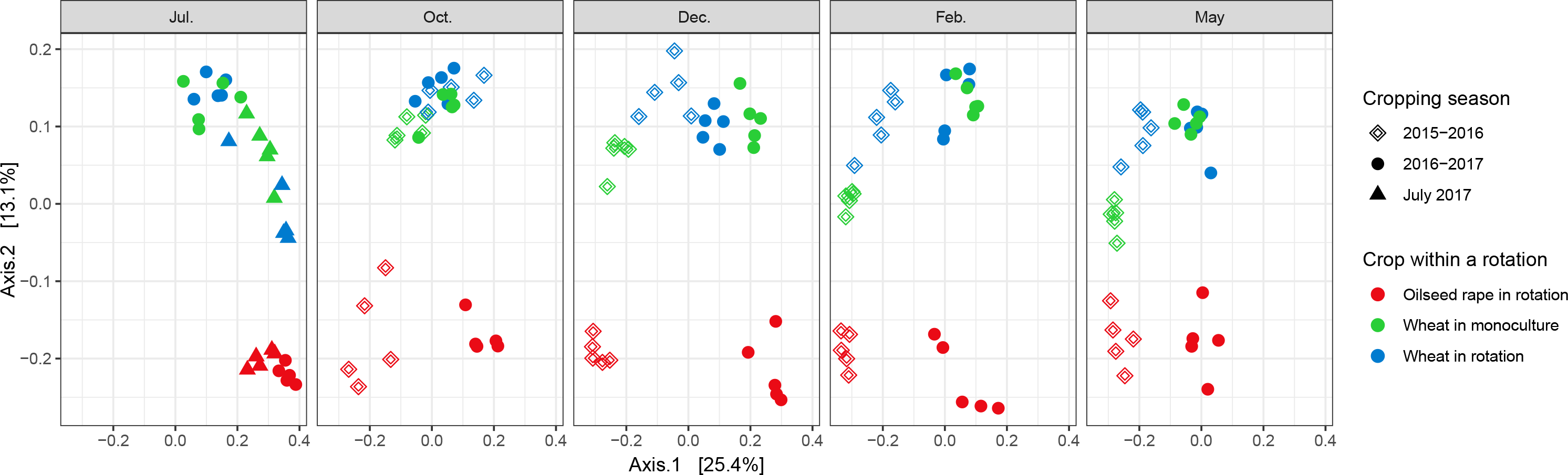
Structure of the bacterial communities present in oilseed rape and wheat residues visualized by incorporating phylogenetic distances using the UniFrac distance matrix. MDS were performed on the overall dataset and faceted according to the sampling period. Each point represents one sample corresponding to a cropping season (shape: 2015-2016; 2016-2017; 2017-2018) and crop within a rotation (colour: oilseed rape in rotation, i.e. in WOW and OWO; wheat monoculture, i.e. in WWW; wheat in rotation, i.e. in WOW and OWO).

**Figure S3.**
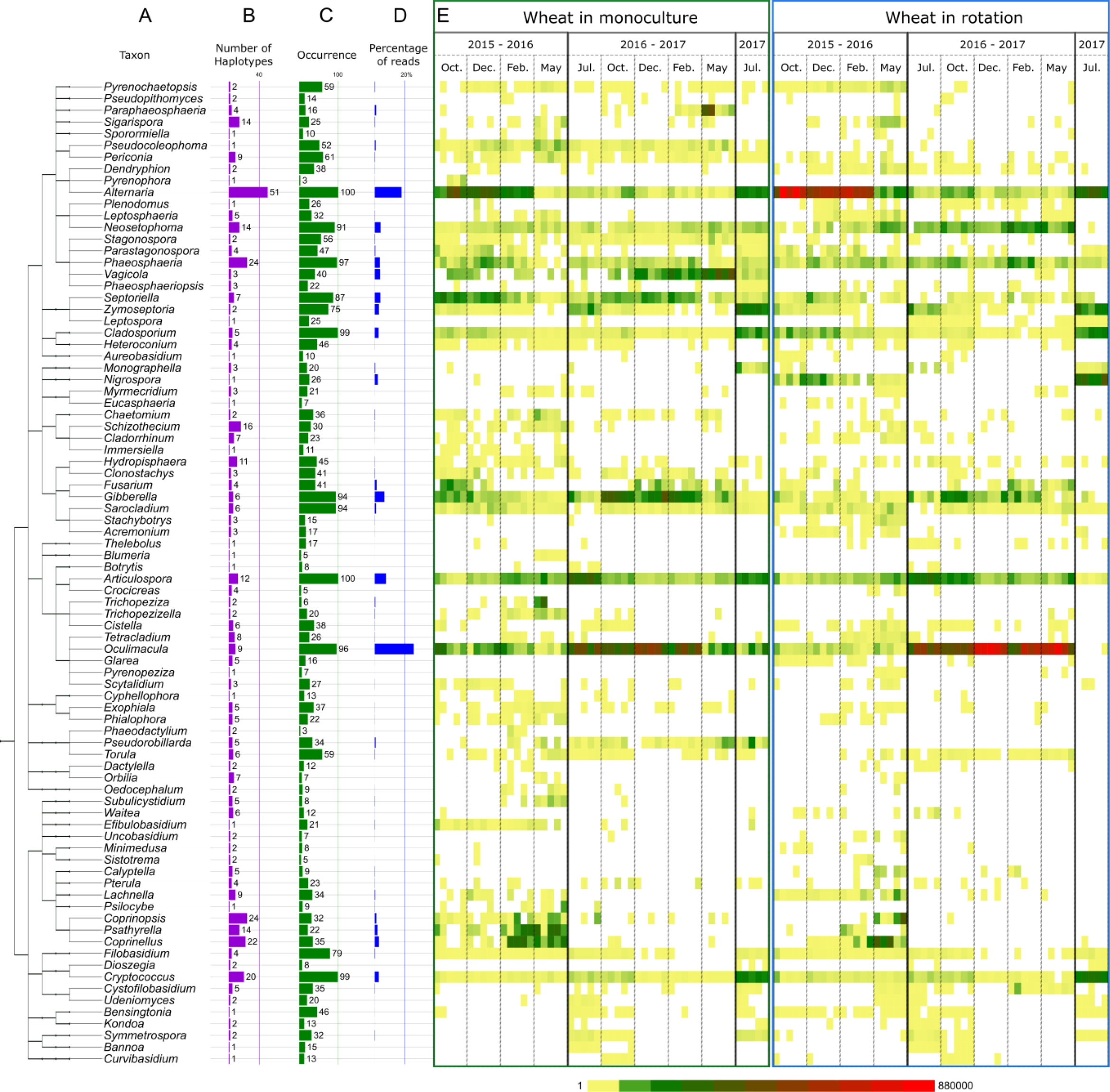
Distribution of the most prevalent fungal genera detected in wheat residues. (**A**) Cladogram of the most prevalent genera. Genera were filtered according to their occurrence (at least three times in the five sampling points for each “crop within a rotation * cropping season * sampling period” combination). Unclassified genera were removed from the tree. (**B**) Number of ASV of each genus. Occurrence of each ASV in the 100 samples of wheat residues. (**D**) Percentage of reads for each genus. (**E**) Distribution of each genus in the five samples per date (increasing number of reads shown on a scale running from yellow to red).

**Figure S4.**
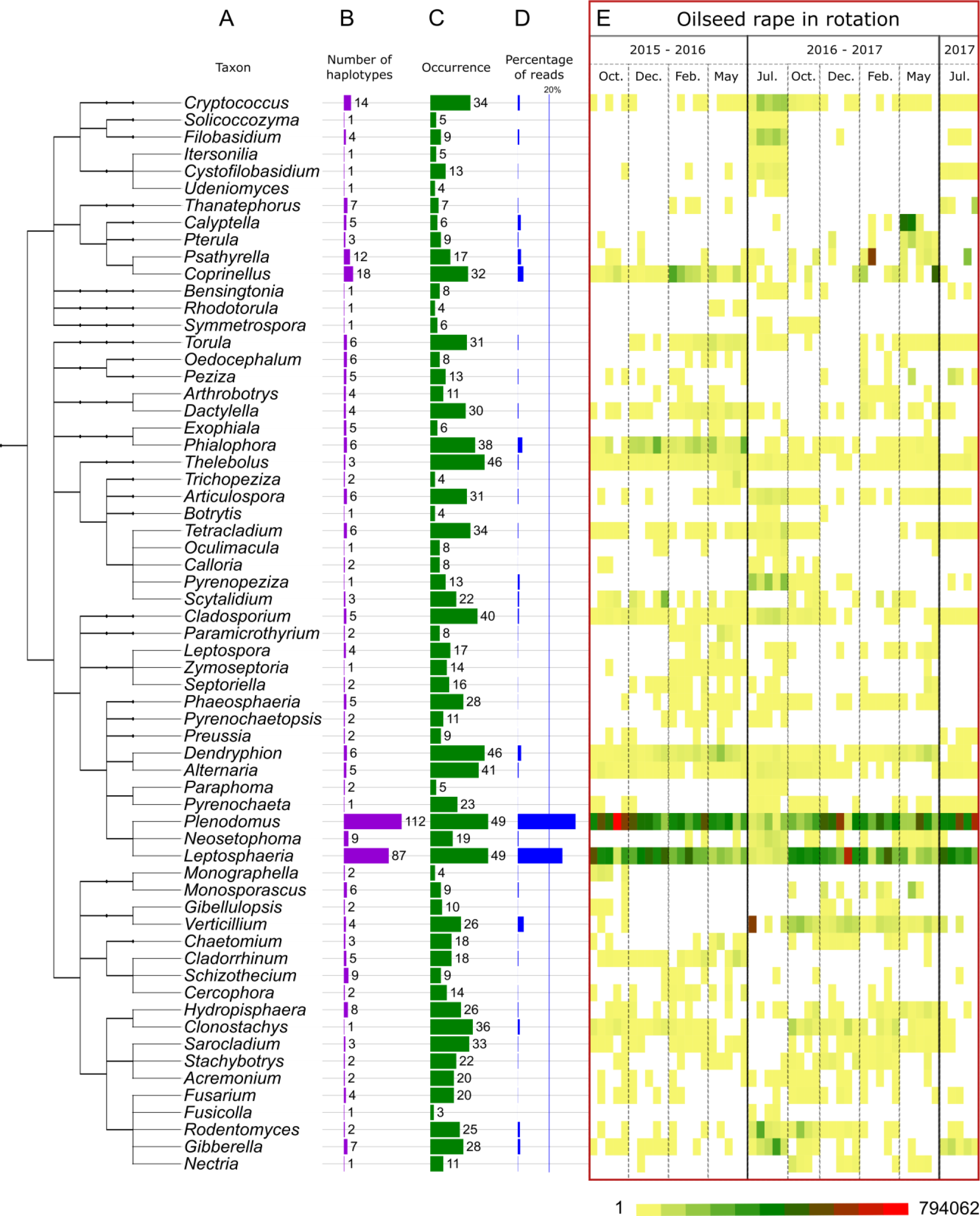
Distribution of the most prevalent fungal genera detected in oilseed rape residues. (**A**) Cladogram of the most prevalent genera. Genera were filtered according to their occurrence (at least three times in the five sampling points for each “crop within a rotation * cropping season * sampling period”) combination. Unclassified genera were removed from the tree. (**B**) Number of ASV for each genus. (**C**) Occurrence of each ASV in the 50 samples of oilseed rape residues. (**D**) Percentage of reads for each genus. (**E**) Distribution of each genus in the five samples per date (increasing number of reads shown on a scale from yellow to red).

**Figure S5.**
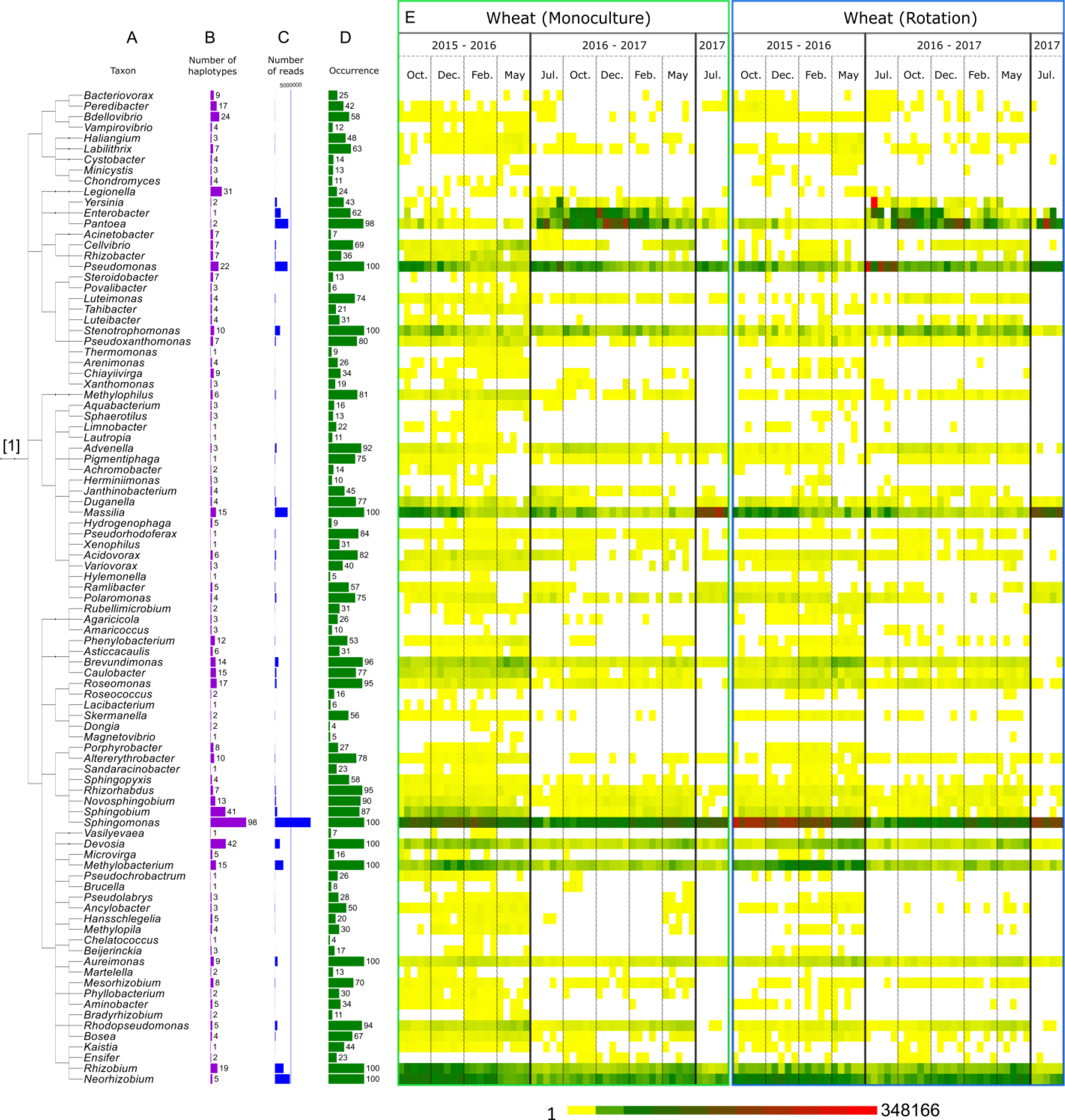
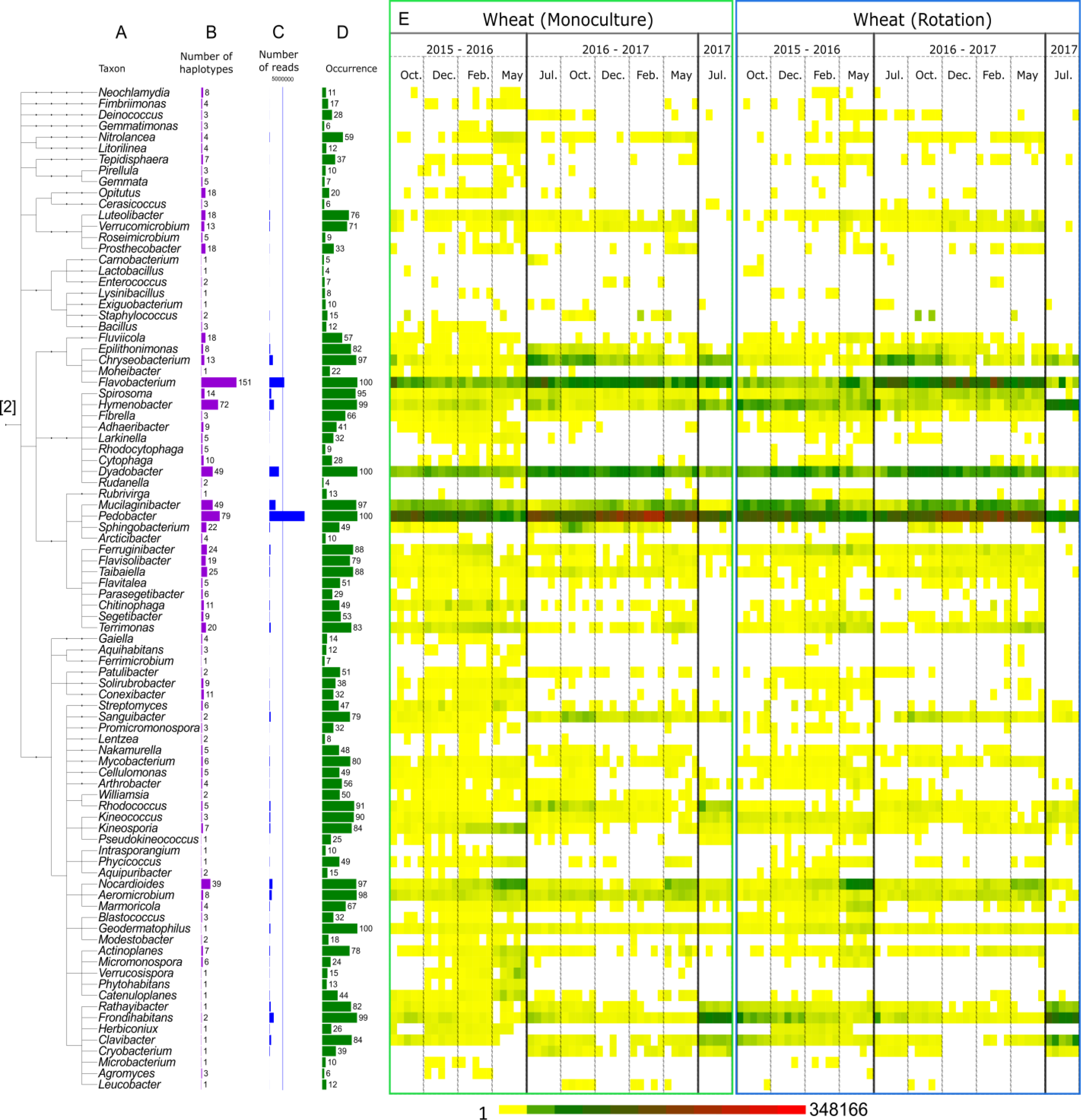
Distribution of the most prevalent bacterial genera detected in wheat residues. **(A)** Cladogram of the most prevalent genera. Genera were filtered according to their occurrence (at least three times in five sampling points for each “crop within rotation * cropping season * sampling period” combination). Unclassified genera were removed from the tree. **(B)** Number of ASV for each genus. **(C)** Occurrence of each ASV in the 100 samples of wheat residues. **(D)** Number of reads for each genus. **(E)** Distribution of each genus in the five samples per date (increasing numbers of reads on a scale running from yellow to red). Due to the number of genera, the plot is separated in [1] proteo-and [2] other bacteria.

**Figure S6.**
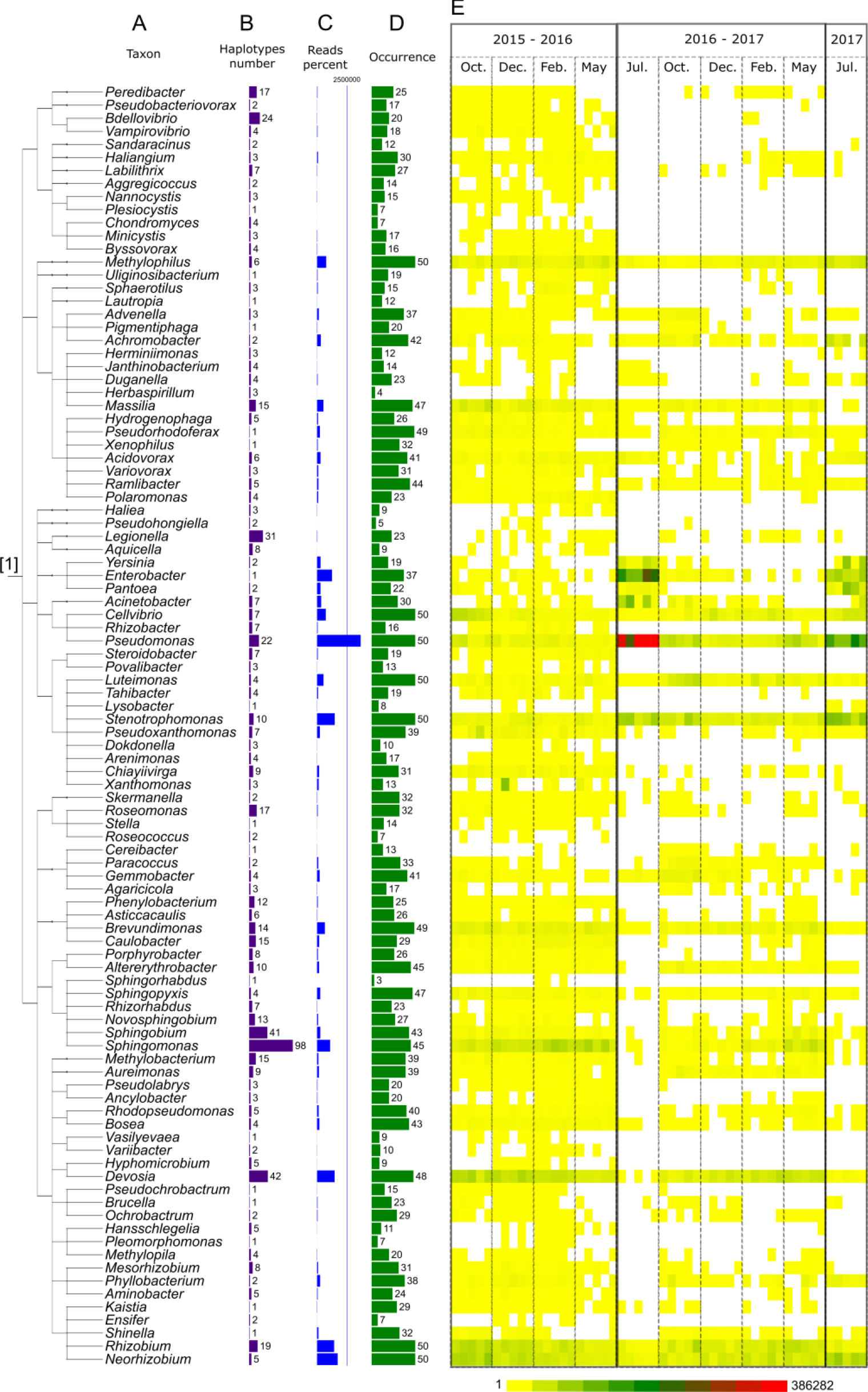
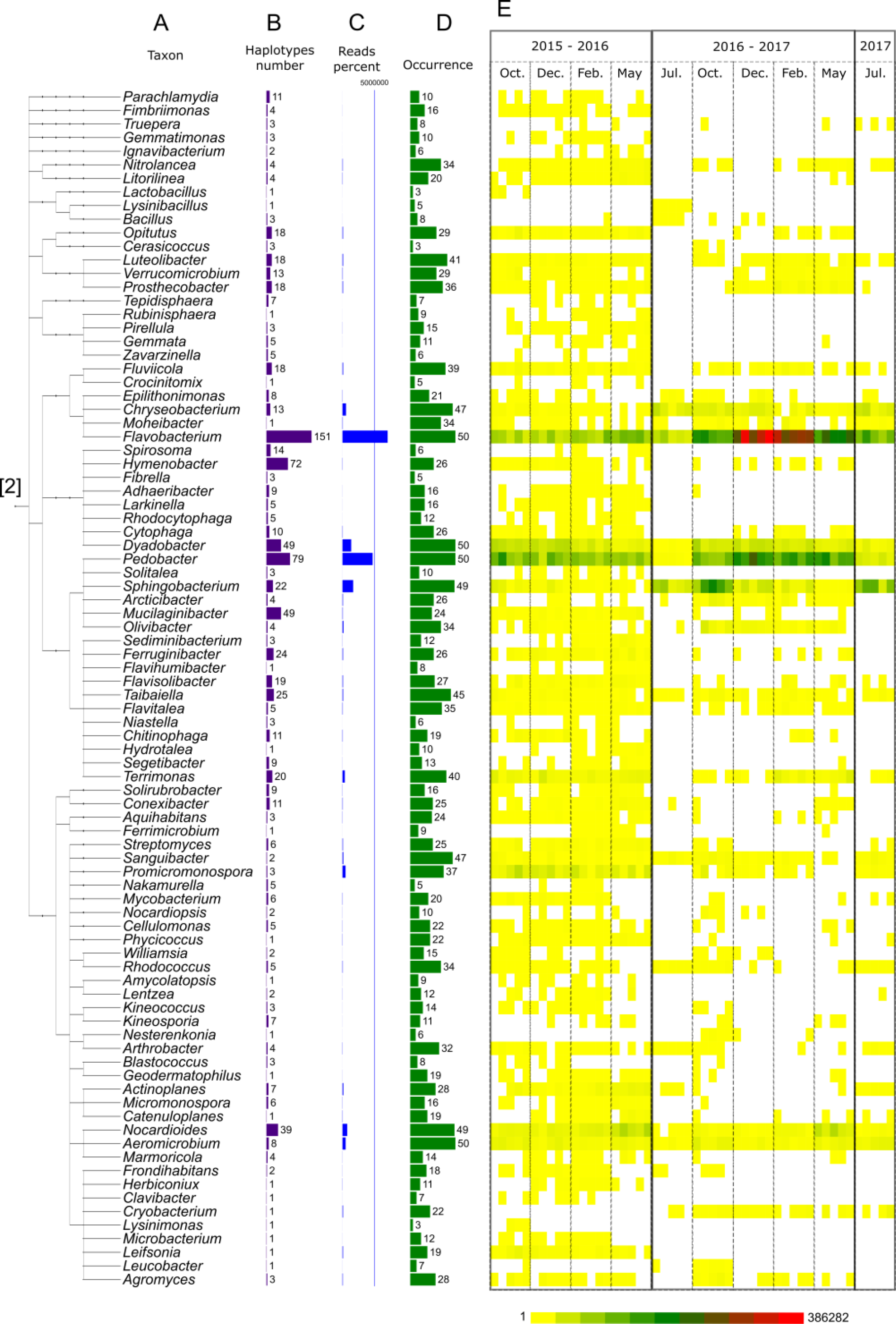
Distribution of the most prevalent bacterial genera detected in oilseed rape residues. **(A)** Cladogram of most prevalent genera. Genera were filtered according to their occurrence (at least three times in five sampling points for each “crop within rotation * cropping season * sampling period” combination). Unclassified genera were removed from the tree. **(B)** Number of ASV for each genus. **(C)** Occurrence of each ASV in the 49 samples of oilseed rape residues. **(D)** Number of reads for each genus. (E) Distribution of each genus in the five samples per date (increasing number of reads shown on a scale running from yellow to red). Due to the number of genera, the plot is separated in [1] proteo-and [2] other bacteria.

**Figure S7.**
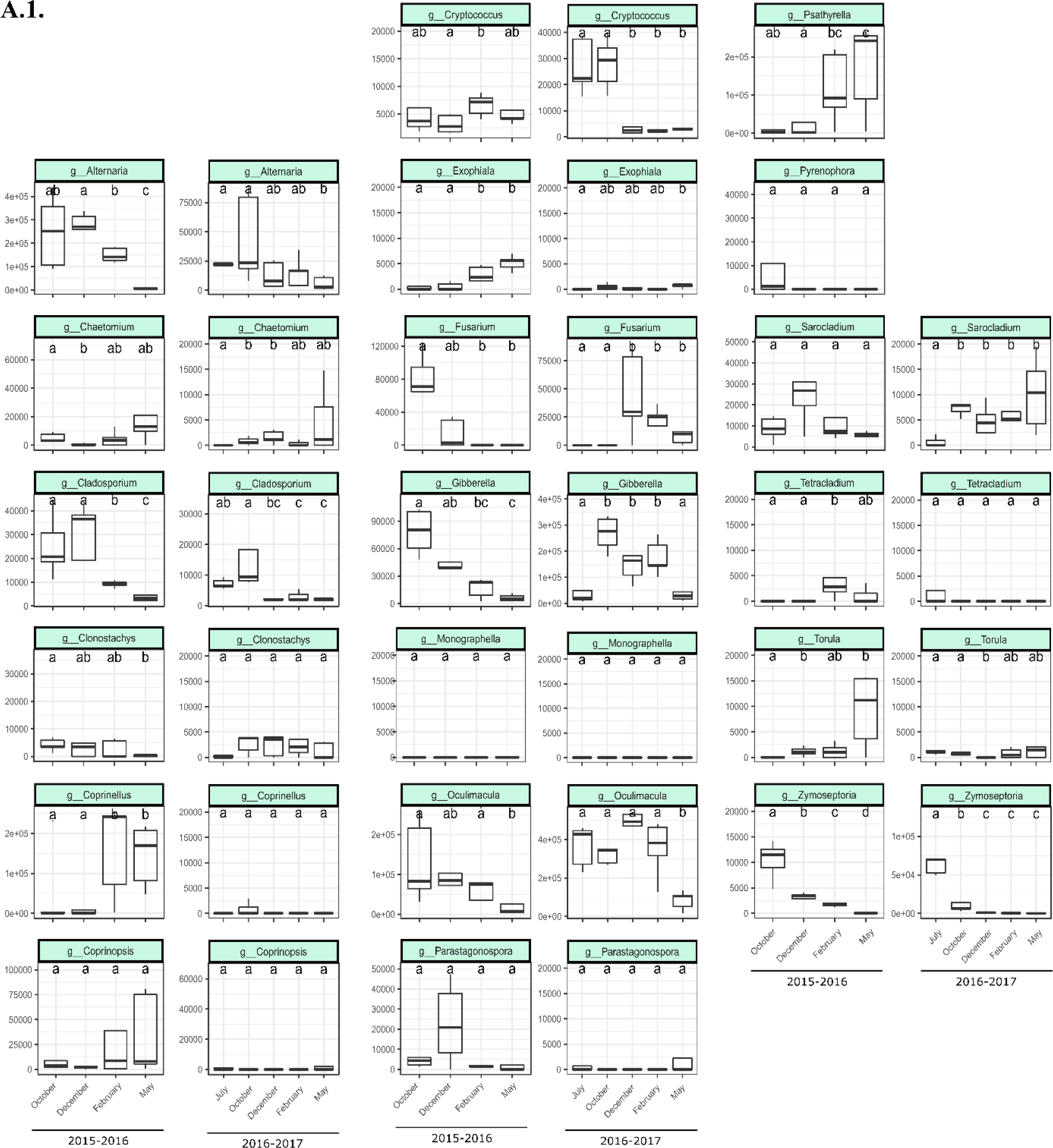
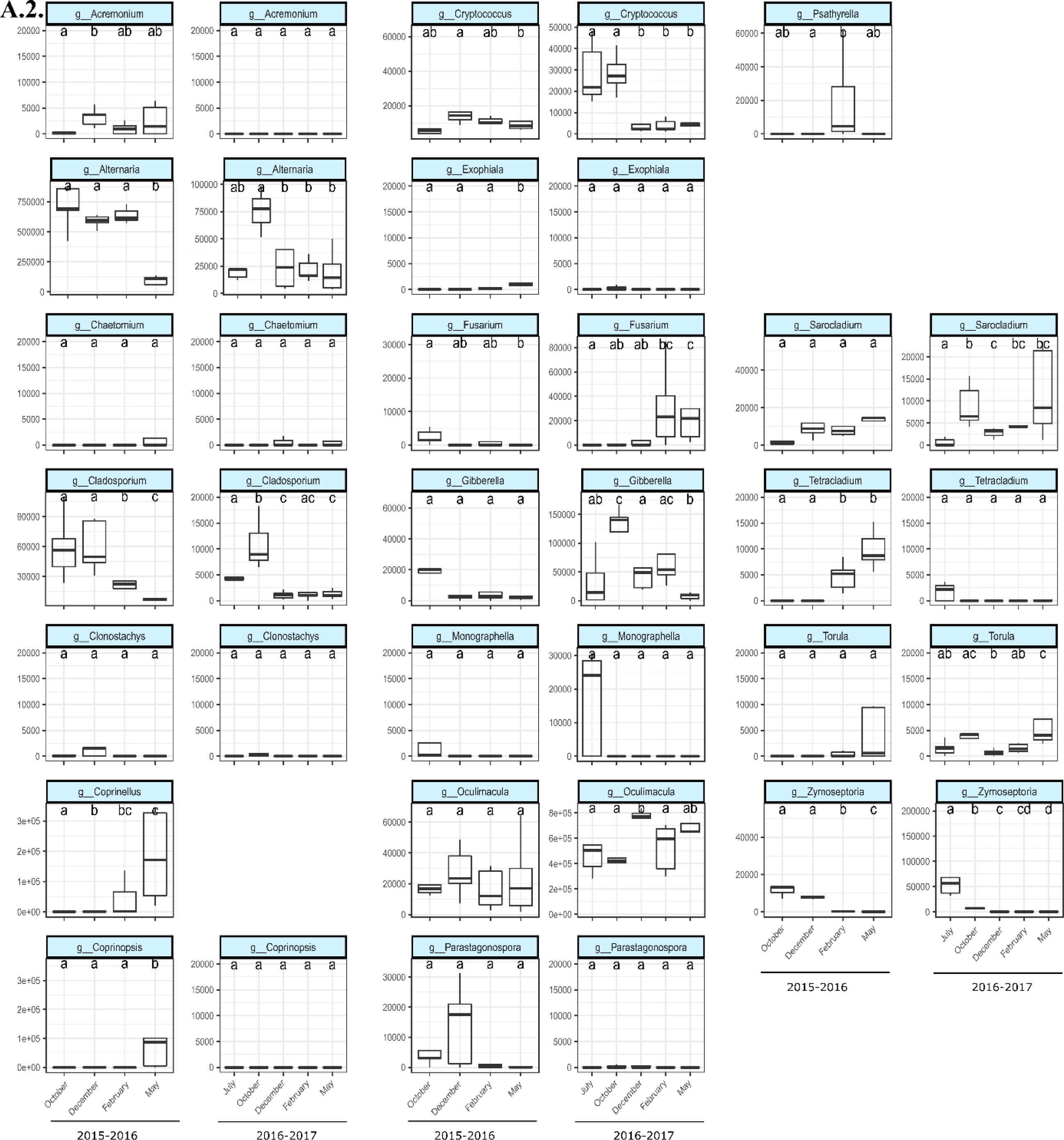
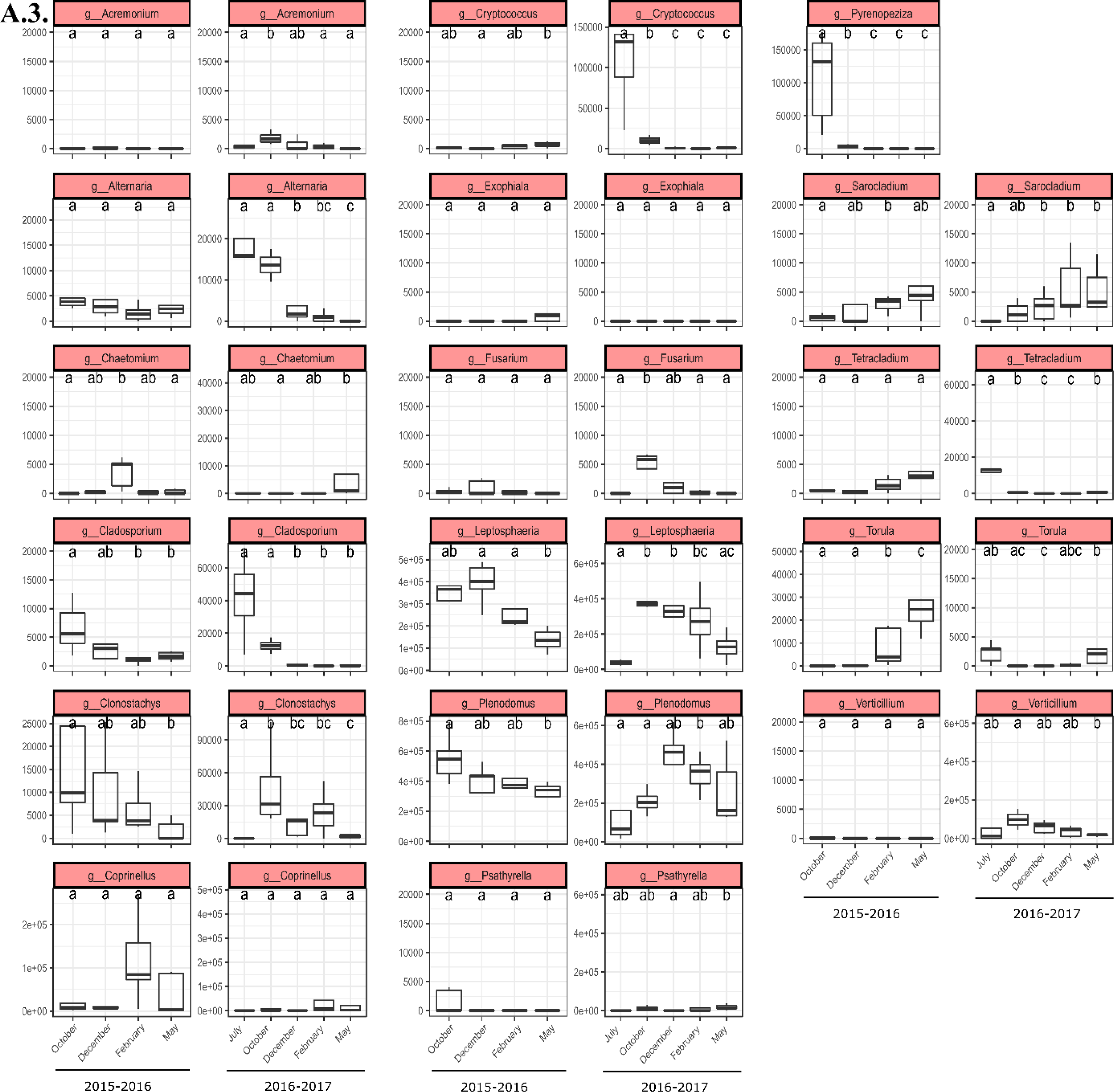
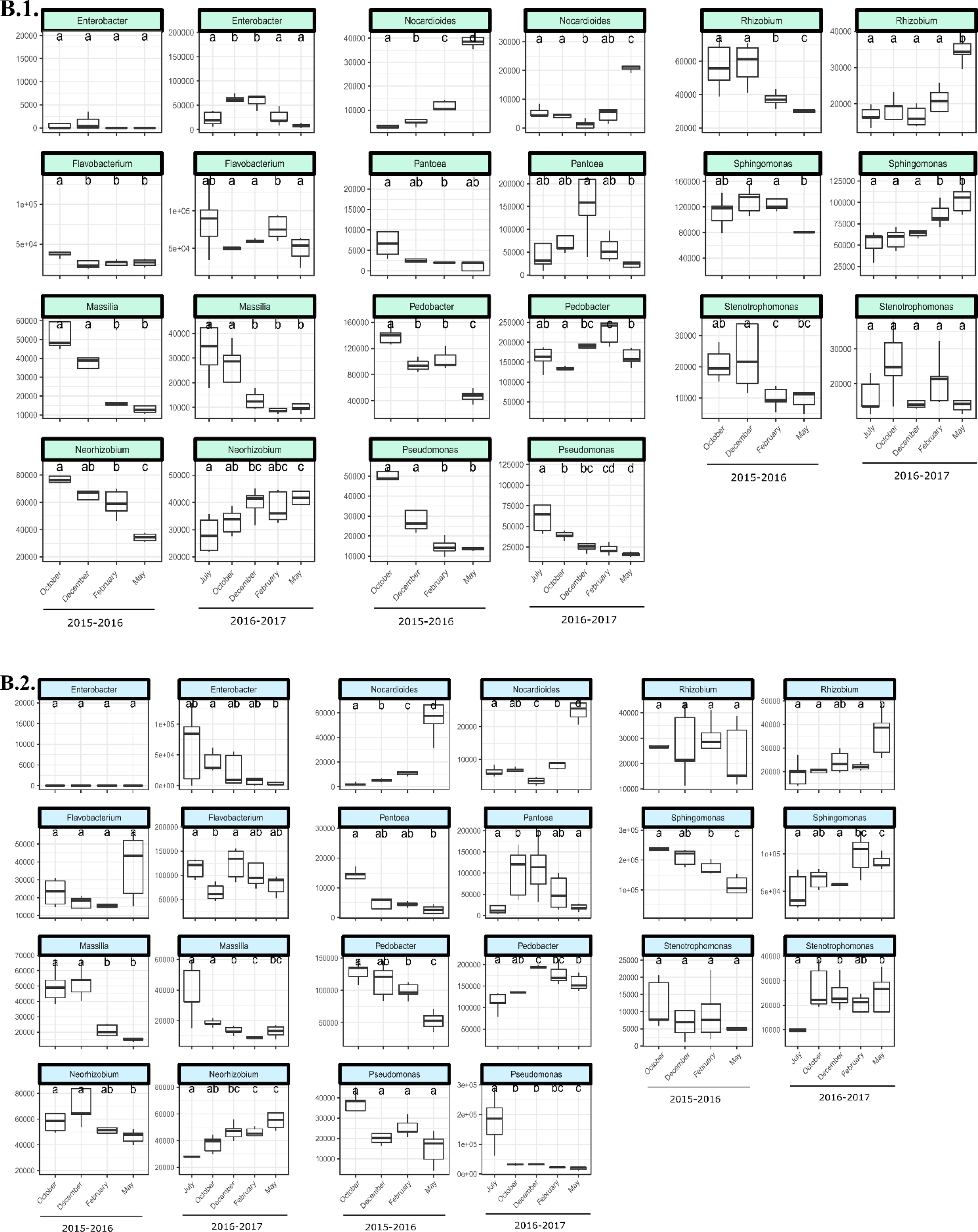
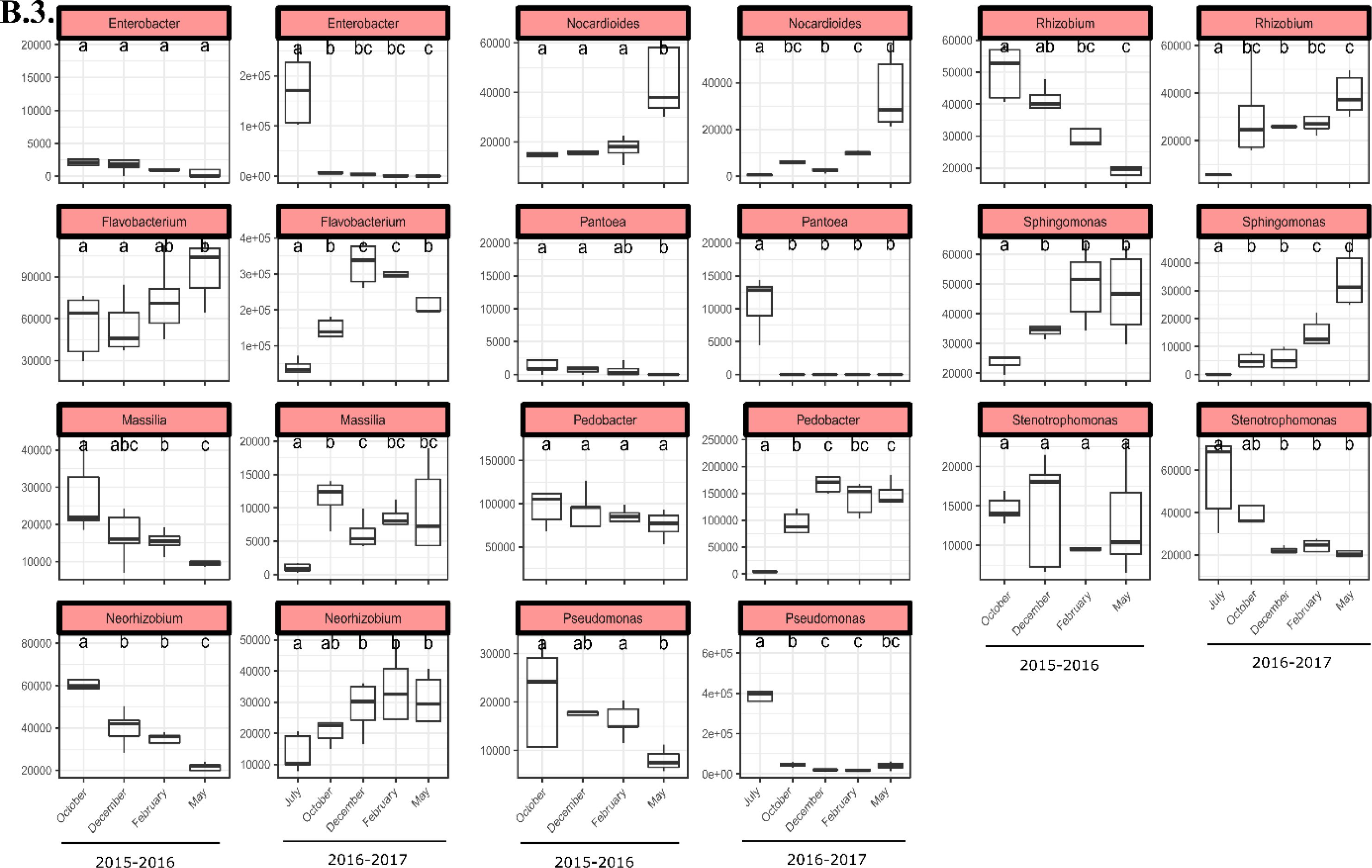
Seasonal shift in the relative abundance of a selection of fungal **(A)** and bacterial **(B)** genera present on wheat and oilseed rape residues according to the system (wheat monoculture [1], wheat in rotation [2], oilseed rape in rotation [3]) and the cropping season (2015-2016, 2016-2017). Due to the plant impact (wheat and oilseed rape) in the fungal community, the fungal genera used here as examples are different for the two plants, unlike the case of the bacterial community. Each box represents the distribution of genera relative abundance for the five sampling points. Samples not sharing letters are significantly different (Wilcoxon tests between sampling periods).

**Table S1.** Soil texture of the three plots (WWW, OWO and WOW).

|  | WWW, OWO | WOW |
| --- | --- | --- |
| Clay (%) | 27.4 | 18.2 |
| Silt (%) | 53.2 | 61.2 |
| Sand (%) | 18.8 | 20.4 |

**Table S2.**
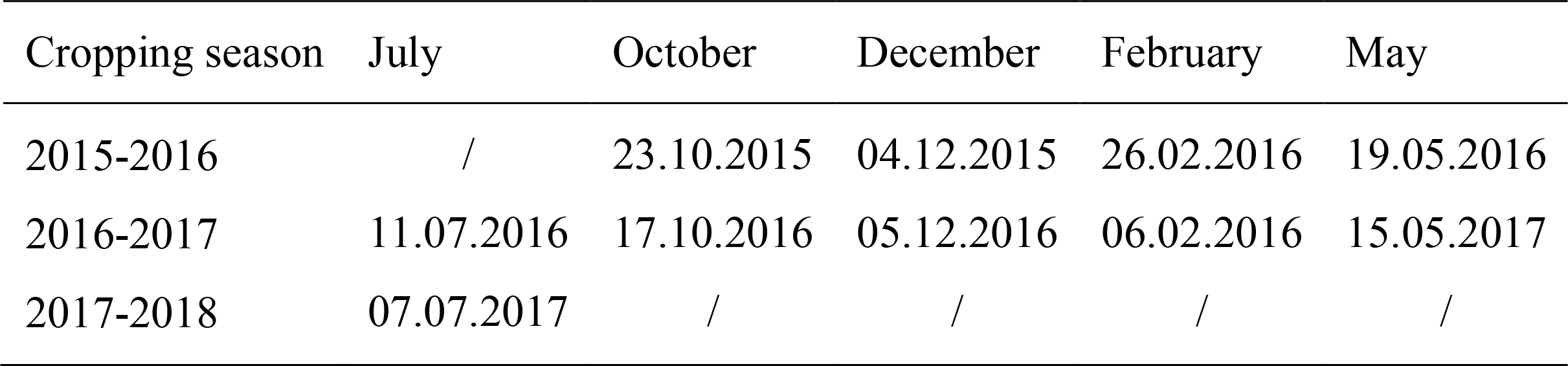
Sampling dates of wheat and oilseed rape plants (July) and residues (October, December, February, and May) for each cropping season.

**Table S3.** Total number of reads and percentage (in brackets) remaining after ASV filtering.

|  | After DADA2 | After replicate suppression | After taxon suppression |
| --- | --- | --- | --- |
| Bacterial reads | 14,287,970 | 13,509,461 (94.6%) | 13,228,976 (92.6%) |
| Fungal reads | 9,898,487 | 9,753,628 (98.5%) | 9,628,995 (97.3%) |
| Bacterial haplotypes | 19,235 | 2,905 | 2,726 |
| Fungal haplotypes | 3,587 | 1,241 | 1,189 |

**Table S4.** Summary of meteorological data (temperature, rainfall) for the INRA Grignon experimental station (Yvelines, France), obtained from the CLIMATIK INRA database (https://intranet.inra.fr/climatik_v2/) from July 1^st^ to May 31^st^ of the following year, for the cropping seasons 2015-2016 and 2016-2017.

|  | 10-day mean temperature (°C) |  | 10-day cumulative rainfall (mm) |  |
| --- | --- | --- | --- | --- |
|  | 2015-2016 | 2016-2017 | 2015-2016 | 2016-2017 |
| Mean | 11.2 | 10.8 | 22.6 | 12.3 |
| Minimum | 2.0 | 0.9 | 0 | 0 |
| Maximum | 21.8 | 21.4 | 131 | 55 |

**Table S5.** Plant effect (wheat vs. oilseed rape) on community dispersion. This effect was tested by applying the adonis function of the vegan R package to the Bray-Curtis dissimilarity index (PERMANOVA). *P*-values (not shown) were all < 0.02.

|  | Fungi |  |  |  | Bacteria |  |  |  |
| --- | --- | --- | --- | --- | --- | --- | --- | --- |
|  | All | 2015-2016 | 2016-2017 | 2017 | All | 2015-2016 | 2016-2017 | 2017 |
| July | 0.372 | / | 0.611 | 0.755 | 0.423 | / | 0.540 | 0.696 |
| October | 0.495 | 0.612 | 0.755 | / | 0.367 | 0.520 | 0.659 | / |
| December | 0.486 | 0.688 | 0.691 | / | 0.370 | 0.573 | 0.641 | / |
| February | 0.429 | 0.541 | 0.651 | / | 0.409 | 0.643 | 0.611 | / |
| May | 0.273 | 0.337 | 0.401 | / | 0.315 | 0.435 | 0.508 | / |

**Table S6.** Decomposition of dissimilarity due to temporal changes in fungal (F) and bacterial (B) community composition. Total dissimilarity is broken down into turnover (replacement of ASV) and nestedness (gain or loss of ASV).

| Crop within a rotation | Season | Sampling period compared | Total dissimilarity |  | Turnover |  | Nestedness |  |
| --- | --- | --- | --- | --- | --- | --- | --- | --- |
|  |  |  | F | B | F | B | F | B |
| Oilseed rape | 2015-2016 | Oct. - Dec. | 0.622 | 0.318 | 0.618 | 0.219 | 0.005 | 0.099 |
| Oilseed rape | 2015-2016 | Dec. - Feb. | 0.650 | 0.321 | 0.577 | 0.290 | 0.073 | 0.031 |
| Oilseed rape | 2015-2016 | Feb. - May | 0.591 | 0.390 | 0.565 | 0.202 | 0.027 | 0.188 |
| Oilseed rape | 2016-2017 | Jul. - Oct. | 0.652 | 0.554 | 0.648 | 0.250 | 0.004 | 0.304 |
| Oilseed rape | 2016-2017 | Oct. - Dec. | 0.620 | 0.353 | 0.549 | 0.217 | 0.071 | 0.136 |
| Oilseed rape | 2016-2017 | Dec. - Feb. | 0.585 | 0.353 | 0.516 | 0.276 | 0.068 | 0.077 |
| Oilseed rape | 2016-2017 | Feb. - May | 0.529 | 0.384 | 0.529 | 0.342 | 0.000 | 0.042 |
| Wheat in monoculture | 2015-2016 | Oct. - Dec. | 0.427 | 0.330 | 0.425 | 0.142 | 0.002 | 0.188 |
| Wheat in monoculture | 2015-2016 | Dec. - Feb. | 0.444 | 0.294 | 0.416 | 0.190 | 0.028 | 0.104 |
| Wheat in monoculture | 2015-2016 | Feb. - May | 0.444 | 0.458 | 0.424 | 0.255 | 0.020 | 0.203 |
| Wheat in monoculture | 2016-2017 | Jul. - Oct. | 0.438 | 0.346 | 0.424 | 0.300 | 0.014 | 0.046 |
| Wheat in monoculture | 2016-2017 | Oct. - Dec. | 0.463 | 0.330 | 0.257 | 0.113 | 0.207 | 0.217 |
| Wheat in monoculture | 2016-2017 | Dec. - Feb. | 0.386 | 0.248 | 0.311 | 0.200 | 0.075 | 0.048 |
| Wheat in monoculture | 2016-2017 | Feb. - May | 0.344 | 0.332 | 0.341 | 0.213 | 0.004 | 0.120 |
| Wheat in rotation | 2015-2016 | Oct. - Dec. | 0.425 | 0.317 | 0.409 | 0.157 | 0.016 | 0.160 |
| Wheat in rotation | 2015-2016 | Dec. - Feb. | 0.472 | 0.266 | 0.370 | 0.185 | 0.102 | 0.081 |
| Wheat in rotation | 2015-2016 | Feb. - May | 0.505 | 0.347 | 0.432 | 0.311 | 0.073 | 0.035 |
| Wheat in rotation | 2016-2017 | Jul. - Oct. | 0.498 | 0.313 | 0.427 | 0.272 | 0.071 | 0.041 |
| Wheat in rotation | 2016-2017 | Oct. - Dec. | 0.541 | 0.287 | 0.292 | 0.214 | 0.249 | 0.073 |
| Wheat in rotation | 2016-2017 | Dec. - Feb. | 0.350 | 0.284 | 0.292 | 0.284 | 0.059 | 0.000 |
| Wheat in rotation | 2016-2017 | Feb. - May | 0.424 | 0.334 | 0.329 | 0.287 | 0.095 | 0.047 |
| Mean |  |  | 0.496 | 0.341 | 0.436 | 0.234 | 0.060 | 0.107 |

